# Butyrate and propionate are microbial danger signals that activate the NLRP3-inflammasome in human macrophages in the presence of TLR stimulation

**DOI:** 10.1101/2024.03.26.586782

**Authors:** Wei Wang, Alesya Dernst, Bianca Martin, Lucia Lorenzi, Maria Cadefau, Kshiti Phulphagar, Antonia Wagener, Christina Budden, Neil Stair, Theresa Wagner, Harald Färber, Andreas Jaensch, Rainer Stahl, Fraser Duthie, Susanne V. Schmidt, Rebecca C. Coll, Felix Meissner, Sergi Cuartero, Matthew S.J. Mangan, Eicke Latz

**Author notes:** Correspondence should be addressed to: Eicke Latz. These authors contributed equally to the project (Wei Wang, Alesya Dernst). These authors contributed equally to the project (Matthew S.J. Mangan, Eicke Latz).

## Abstract

Short chain fatty acids (SCFAs) are immunomodulatory compounds produced by the microbiome through fermentation of dietary fibre. Although they are generally considered beneficial for gut health, patients suffering from inflammatory bowel disease (IBD) have shown poor tolerance to fibre-rich diets, suggesting that SCFAs may have contrary effects under inflammatory conditions. To investigate this, we examined the effect of SCFAs on human macrophages in the presence of toll-like receptor agonists. In contrast to their anti-inflammatory effects under steady state conditions, we observed that the SCFAs butyrate and propionate triggered the activation of the NLRP3 inflammasome when added in conjunction with TLR agonists. Mechanistically, butyrate and propionate activated NLRP3 by inhibiting HDACs 1-3 and 10, leading to an uneven distribution of histone hyperacetylation that resulted in alterations in the transcriptome. Specifically, there was a lack of hyperacetylation at the loci of the *CFLAR* and *IL10* genes, two important inhibitors of NLRP3 inflammasome activation. The concurrent loss of transcription and protein expression of cFLIP and IL-10 enabled caspase-8-dependent NLRP3-inflammasome activation. SCFA-driven NLRP3 activation did not require potassium efflux and did not result in cell death but rather triggered hyperactivation and IL-1β release. Our findings demonstrate that butyrate and propionate are bacterially-derived, viability-dependent danger signals (vita-PAMPs) that regulate NLRP3 inflammasome activation through epigenetic modulation of the inflammatory response.

**Summary:** Under inflammatory conditions, SCFAs are bacterially-derived, viability-dependent danger signals that, through HDAC inhibition and epigenetic modification, prevent expression of the anti-cell death gene cFLIP to trigger activation of the NLRP3 inflammasome.

## Introduction

Short chain fatty acids (SCFAs), including acetate, propionate and butyrate, are generated at high concentrations by the microbiota through fermentation of dietary fibre ^1^. This occurs mostly in the intestine, where, in the ileum and descending colon concentrations can range from 10 to 90 mM at a ratio of 3:1:1 acetate:propionate:butyrate. Additionally, SCFA production takes place in the periodontal pockets in the mouth ^1,2^. Under steady state conditions, SCFAs play a crucial role in maintaining gut homeostasis. They contribute to this balance by both supporting the integrity of the intestinal epithelial barrier and modulating the function of immune cells. SCFAs mediate these effects by activation of cell surface receptors for free fatty acids, G protein-coupled receptors (GPCRs) and olfactory receptors. Additionally, they diffuse into the cell, where they can inhibit histone deacetylase (HDAC) activity or serve as a fuel source ^3–5^. Through these mechanisms SCFAs regulate the immune system to limit inflammation and promote immunity. One notable example is their effect on intestinal macrophages, where butyrate, through epigenetic modification, triggers differentiation of monocytes to macrophages and imprints an anti-inflammatory, anti-microbial transcriptional program ^6^.

Due to their action as immune modulators, SCFAs have been studied as a potential therapy for intestinal inflammatory disorders, including inflammatory bowel disease (IBD) ^7^. IBD is characterized by a breakdown of the intestinal epithelial cell barrier, exposing underlying immune cells to microbial products. It is driven by a combination of genetic and environmental factors alongside immune dysfunction ^7^. Given the complexity IBD development, treatment has primarily focused on alleviating symptoms by reducing the inflammatory reaction. One proposed treatment for IBD involves fibre-derived SCFAs. However, despite expectations, it remains unclear whether SCFAs are beneficial or detrimental for IBD, as fibre, which enables SCFA production, is poorly tolerated by IBD patients in some cases ^8,9^. This intolerance is partially attributed to the type of fibre ingested ^10^, but importantly diets that increase the level of SCFAs do not always have beneficial effects ^11^. This suggests that fibre, and, consequently, SCFAs may have different actions depending on the context in which they are generated.

The role of inflammation in IBD has suggested that drugs targeting inflammatory cytokines involved in the pathophysiology of IBD would have therapeutic potential. However, so far, drugs targeting TNFα, IL-12 or IL-6 have been unable to completely revert the IBD phenotype ^12–14^. Subsequently, the NLRP3 inflammasome, which triggers secretion of IL-1β and IL-18, has emerged as a new potential target in the treatment of IBD. Notably, people harbouring hyperactivating NLRP3 mutations have increased risk of ulcerative colitis and IBD ^15,16^. This is less clear in two murine models of colitis, dextran sulfate sodium (DSS) or 2,4,6-trinitrobenzenesulfonic acid (TNBS), where NLRP3 has been suggested to be both detrimental or protective in the disease ^17–20^. Furthermore, the ablation of caspase-1 rescued spontaneous colitis that occurs in the IL-10-deficient murine model ^21^. The variations in the results likely reflects the complex aetiology of IBD and differences arising from disparities in genetic background, housing conditions, and microbiota composition. Perhaps the most relevant result then is that CP-456,773 (also called CRID3 or MCC950) mitigated DSS colitis in littermate control with the same genetic background ^17^, indicating that NLRP3 is a relevant target for treatment of IBD.

In this study we address this question by examining the effect of SCFAs on the LPS- mediated inflammatory response in human GM-CSF derived macrophages to understand the underlying mechanisms of SCFAs in an inflammatory context. Surprisingly, we observed that SCFAs, in addition to reducing LPS-mediated cytokine transcription, activated the NLRP3 inflammasome when combined with TLR agonists. Both butyrate and propionate triggered HDAC-inhibition dependent histone hyperacetylation across the genome, altering gene transcription in a HAT-dependent manner. Specifically, HDAC inhibition resulted in a failure to hyperacetylate the loci of two important regulators of NLRP3 inflammasome activation, *CFLAR* and *IL10* loci, leading to a loss of cFLIP and IL-10 expression. The concurrent loss of expression of both of these proteins then enabled caspase-8 dependent NLRP3 inflammasome activation.

## Results

### Butyrate and propionate downregulate the LPS-driven cytokine response but trigger IL-1β release

To investigate the impact of SCFAs on the inflammatory response in human macrophages, we incubated human monocyte-derived macrophages, differentiated with GM-CSF (hMDM), with LPS in the presence or absence of the SCFAs butyrate, propionate or acetate. We measured cytokine secretion from supernatants after a 16-hour incubation using a broad-range cytokine bead array. Compared to LPS in the presence of NaCl, the vehicle control, co-incubation of LPS with SCFAs decreased secretion of most LPS-dependent cytokines, including: IL-10, IL-12p40, IL-6 and TNFα (Fig. 1A). Among the three SCFAs, butyrate exhibited the strongest effect, noticeable already at 1 mM, while propionate exhibited a moderate effect and acetate had almost no effect. Notably, and in contrast to the majority of the assessed cytokines, butyrate and propionate triggered release of IL-1β, a key pro-inflammatory cytokine, in the presence of LPS (Fig. 1A). This was particularly intriguing since, unlike most measured cytokines, IL-1β release occurs post-translationally, and is often associated with the activation of caspase-1, the effector of the inflammasome complex. To further characterise this phenomenon, we assessed IL-1β release over time and found that the combination of butyrate and LPS triggered IL-1β release as early as 6 h, increasing up to 18 h (Fig. 1B). Maximal secretion of IL-1β occurred at 4 mM butyrate, although substantial IL-1β release was already observed at 1 mM (Fig. S1A). Neither butyrate nor LPS alone triggered IL-1β release (Fig. 1B). Considering that inflammasome activation can lead to pyroptotic cell death, we assessed this by measuring LDH release. Interestingly, LDH was not released into the media (Fig. 1C), indicating that butyrate did not cause lytic cell death.

**Figure 1.**
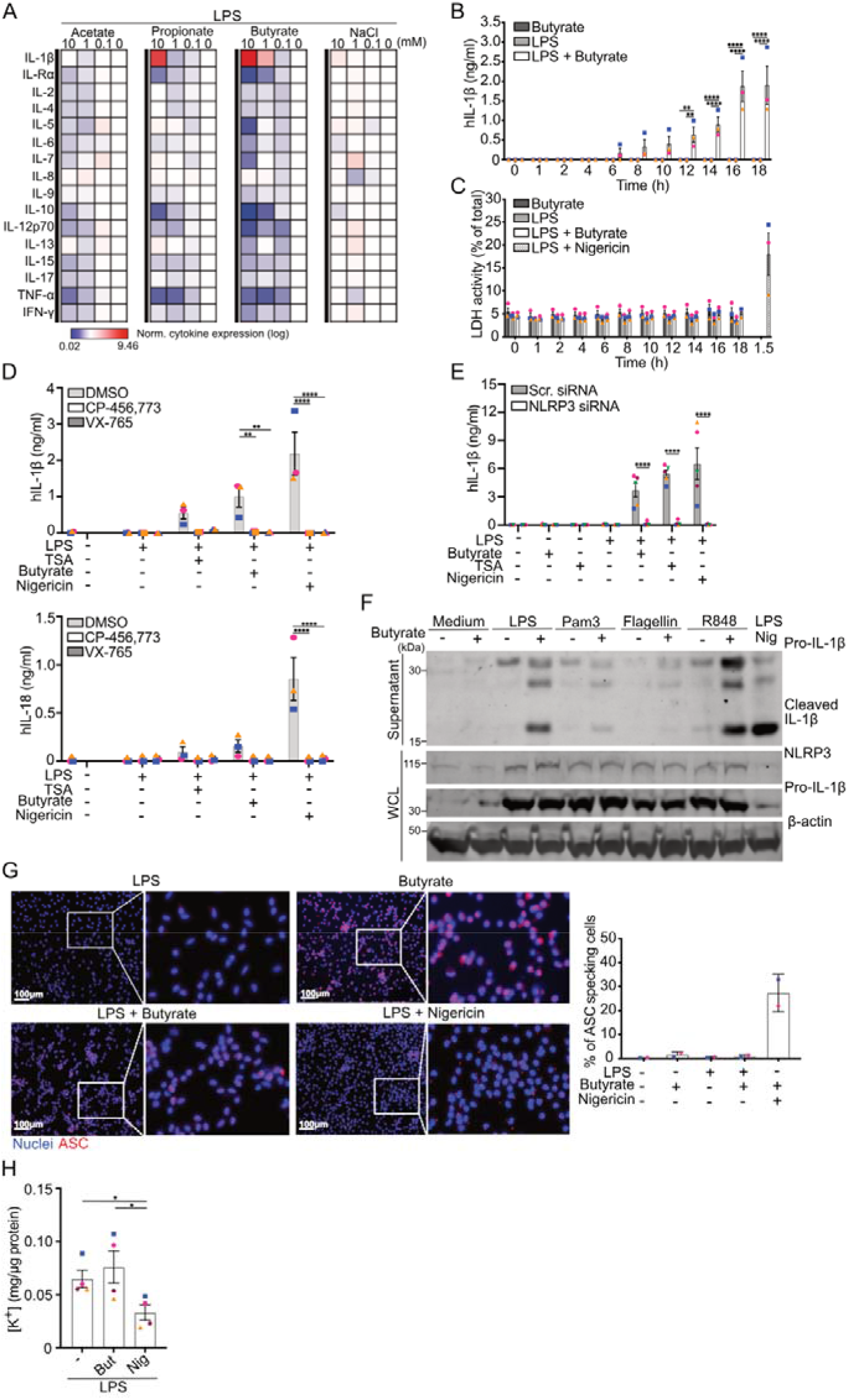
SCFAs decrease LPS-induced cytokine secretion but trigger NLRP3- inflammasome dependent IL-1β release. (A) hMDM were treated with either NaCl, acetate, propionate or butyrate (all 0.1-10 mM) in the presence of LPS (1 ng/ml) for 16 h. Heat map of cytokines measured from cell free supernatants in response to stimulation with LPS (1 ng/ml) +/- acetate, propionate or butyrate and normalised to secretion from cells treated with LPS and NaCl. Mean value of 6 different donors is shown in logarithmic scale. (B) IL-1β or (C) LDH release measured from cell-free supernatants from hMDM treated with LPS (1 ng/ml), butyrate (1 mM) or a combination for the listed times. IL-1β release measured from cell-free supernatants from hMDM treated with butyrate (10 mM) alone or with LPS (1 ng/ml) for 16 h or primed with LPS (1 ng/ml, 3 h) and stimulated with nigericin (10 μM 1.5 h) pre-incubated with or without (D) VX-765 (40 μM) or CP-456,773 (2 μM) for 30 min. Data shown represent 3 (B, C and D lower panel) or 5 (D upper panel) independent experiments. (E) IL-1β release from hMDM electroporated with NLRP3 or scrambled siRNA, then treated with butyrate (10 mM) alone or with LPS (1 ng/ml) for 16 h, TSA (0.5 μM) with LPS (1 ng/ml) for 16 h or nigericin (10 μM 1.5 h). (F) Immunoblot of cell free supernatants or whole cell lysate (WCL) of hMDM treated with butyrate (10 mM) alone or with LPS (1 ng/ml), Pam3CSK4 (10 ng/ml), Flagellin (50 ng/ml) or R848 (250 ng/ml) for 16 h. Representative of 3 independent experiments. (G) Images of LPS-primed hMDM stained for ASC (red) and nuclei (blue) treated with LPS and butyrate (10 mM, 16 h) or LPS and nigericin (10 μM 1.5 h) in the presence of VX-765 (40 μM). Quantitation of ASC specks shown for 4 images per condition. (H) Potassium levels from hMDM treated as in (D) and analysed by ICP-MS. Data shown represents 4 independent experiments. Data are represented as mean +/- SEM, unless otherwise indicated, each dot is representative of one donor in all cases, *p<0.05, **p < 0.01, ***p < 0.001, ****p < 0.0001 by Two-way ANOVA with SΛídák’s multiple comparisons test.

### Butyrate-mediated IL-1**β** release is dependent on the NLRP3 inflammasome in hMDM

A key mechanism driving IL-1β release is activation of an inflammasome complex. Among these complexes, NLRP3 is the most notable, triggering inflammasome formation and caspase-1 activation in response to a diverse range of cell stresses ^22^. Given the effects of butyrate and propionate on the inflammatory response, we first verified that expression of NLRP3 and pro-IL-1β was not changed by incubation with butyrate (Fig. S1B). To investigate whether NLRP3 was required for butyrate-mediated IL-1β release, we incubated hMDM with a caspase-1 inhibitor (VX-765) or an NLRP3 inhibitor (CP-456,773, also known as MCC950 or CRID3) prior to addition of inflammasome stimuli. In addition to butyrate, we stimulated the hMDM with trichostatin A (TSA), a pan-HDAC inhibitor, as both butyrate and propionate, but not acetate, have HDAC inhibitory activity^23^. Both butyrate and TSA triggered IL-1β release, demonstrating that inflammasome activation is likely mediated by HDAC inhibition. Both VX-765 and CP-456,773 were found to inhibit TSA- and butyrate-mediated IL-1β release (Fig. 1D) as well as butyrate-mediated IL-1β cleavage (Fig. S1C), indicating that butyrate-driven IL-1β release is dependent on the NLRP3 inflammasome. Both butyrate and TSA also mediated IL-18 release, though at a much lower level than IL-1β, and this was blocked by both inhibitors (Fig. 1D), though this was not significant. Nigericin, a known activator of the NLRP3 inflammasome, was similarly inhibited by both compounds, preventing both IL-1β and IL-18 release (Fig. 1D). To confirm the role of NLRP3, we transfected hMDM with siRNA targeting either NLRP3 or a scrambled control. The NLRP3-targeting siRNA reduced NLRP3 expression (Fig. S1D) and prevented IL-1β release in response to butyrate, TSA and nigericin (Fig. 1E). In contrast, the NLRP3-targetting siRNA had no effect on TNFα secretion (Fig. S1E). This was not limited to activation of TLR4, as stimulation of hMDM with Pam3CSK4 (TLR2), flagellin (TLR5) and R848 (TLR7/8) also stimulated IL-1β cleavage and release (Fig. 1F, S1F).

Another hallmark of inflammasome activation is the aggregation of the inflammasome adapter ASC, termed an ASC speck or pyroptosome. hMDM were treated with or without butyrate, with or without LPS, in the presence of VX-765, and assessed for ASC speck formation, using LPS and nigericin as controls. Surprisingly, hMDM treated with LPS and butyrate triggered only a minor number ASC speck formation (Fig. 1G), suggesting these are not required for the butyrate-driven NLRP3 response. This was in contrast to nigericin, where robust ASC speck formation was evident (Fig. 1G).

The absence of ASC speck formation was reminiscent of the alternative pathway of activation observed in human monocytes in response to prolonged exposure to LPS ^24^. Unlike the majority of NLRP3 activators, alternative NLRP3 activation does not trigger potassium efflux. To determine if this was also true for LPS and butyrate-driven NLRP3 activation, we measured potassium levels using inductively coupled plasma mass spectrometry (ICPMS) in the hMDM following stimulation with LPS, LPS and butyrate or LPS and nigericin (an activator that triggers potassium efflux). We determined that LPS and butyrate did not cause a decrease in the cellular potassium concentration compared to the hMDM treated with LPS alone (Fig. 1H), while there was a marked decrease in response to nigericin (Fig. 1H). This indicates that butyrate-driven NLRP3 activation does not require potassium efflux.

### Inhibition of HDACs 1-3 and 10 is sufficient to trigger NLRP3-dependent IL-1**β** release

SCFAs mediate many of their functions through the inhibition of HDAC activity ^25^. The propensity to activate NLRP3 correlated with the effectiveness of the respective SCFA to inhibit HDAC activity. Furthermore, TSA, a pan-HDAC inhibitor, also triggered NLRP3-depdendent IL-1β release. This contrasts to a previous study showing that HDAC inhibition triggered NLRP3-independent IL-1β release^26^. We tested whether HDAC inhibition would be sufficient to recapitulate the effects of butyrate by incubating hMDM with LPS and a range of HDAC inhibitors with varying specificities (Fig. 2A). Pan-HDAC inhibitors, which overlap in specificity with butyrate, also triggered NLRP3 activation (Fig. 2B). Notably, inhibitors that targeted a combination of HDACs 1-3 and 10 could all mediate IL-1β release in the presence of LPS (Fig. 2B), but did not perturb secretion of TNFα (Fig. 2C). This indicates that inhibition of HDAC activity enables spontaneous NLRP3 inflammasome activation under inflammatory conditions.

**Figure 2.**
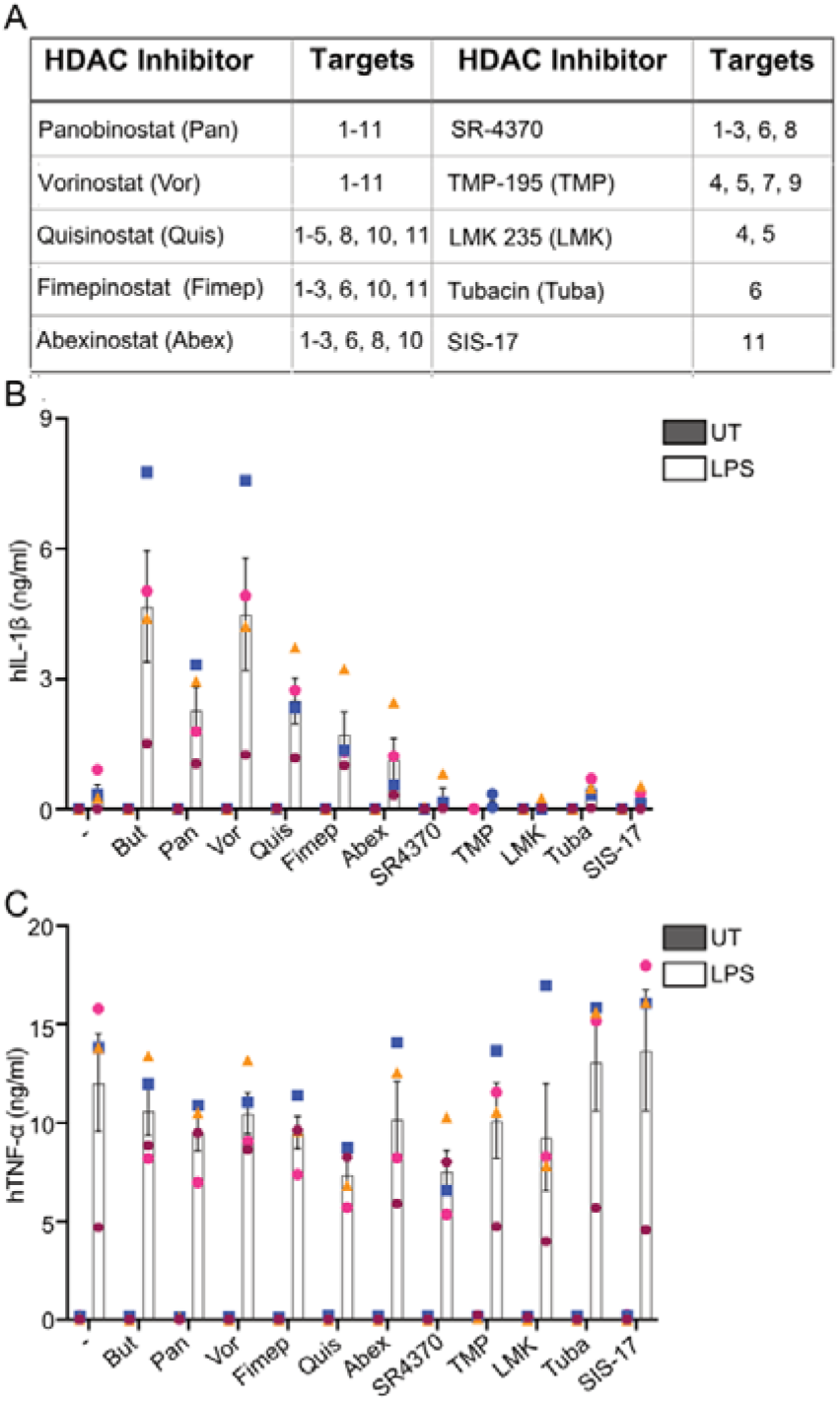
Inhibition of HDACs 1-3 and 10 enables NLRP3 activation in response to TLR stimulation. (A) The concentration of HDAC inhibitors used and their specificities. (B) IL-1β release or (C) TNFα secretion measured from cell-free supernatants from hMDM treated with HDAC inhibitors alone or with LPS (1 ng/ml) for 16 h. Concentrations used were Butyrate (10 mM), Pan (1 μM), Vor (1 μM), Quis (0.2 μM), Fimep (0.1 μM), Abex (0.1 μM), SR4370 (4 μM), TMP (0.3 μM), LMK (20 nM), Tuba (50 nM) and SIS-17 (1 μM). In all cases each dot represents one donor, mean and SEM shown.

### Butyrate and propionate modulate the LPS-driven gene expression profile

Activation of NLRP3 by butyrate and propionate required the inhibition of HDAC activity, which, along with a relatively long incubation period, suggesting that it was occurring through changes in transcription. To assess the effect of SCFAs on the LPS-mediated transcriptome in hMDM, we conducted RNA-seq in hMDM treated with butyrate, propionate or acetate in the presence of LPS for 16 h. NaCl was included as the vehicle alone control, and TSA was included to define the effect of HDAC inhibition on the LPS response.

The principal component analysis demonstrated that LPS accounted for the largest differences between the treatment groups (Fig. 3A). Butyrate and TSA clustered together, suggesting most alterations caused by butyrate are due to its HDAC inhibitory activation (Fig. 3A). Indeed, a comparison of genes changed by butyrate and TSA revealed almost overlapping gene signatures (Fig. S2A). Notably, in addition to LPS-independent effects, both partially reversed the effect of LPS. Analysis of the most changed genes by hierarchical clustering supported this conclusion, as in many cases butyrate and TSA reversed the effect of LPS, either increasing expression of genes decreased by LPS or vice versa (Fig. 3B). Propionate had a similar but weaker effect compared to butyrate and TSA (Fig. 3A), also showing an overlapping signature with butyrate (Fig. S2B) while acetate had minimal effect (Fig. 3A). It was interesting to note that, in spite of histone acetylation generally being positively associated with gene transcription, butyrate, TSA and propionate resulted in both the up- and downregulation of a similar number of genes (Fig. 3C). To investigate this further we performed gene set enrichment analysis (GSEA) using the hallmark gene sets on genes altered by the different treatments. Notably, the GSEA for butyrate and TSA, and to a lesser extent propionate, were enriched for almost exactly the same terms associated with downregulated genes, including inflammatory cytokine response, the interferon response or the TNFα pathway (Fig. 3D). In contrast, the few terms that were significantly enriched in upregulated genes were not consistently identified among the three treatments, indicating that no specific pathways were activated by SCFAs (Fig. 3D). The correlation between the enriched pathways from the SCFAs and TSA suggested that the primary effect of SCFAs on the LPS-driven gene signature was due to HDAC inhibition.

**Figure 3.**
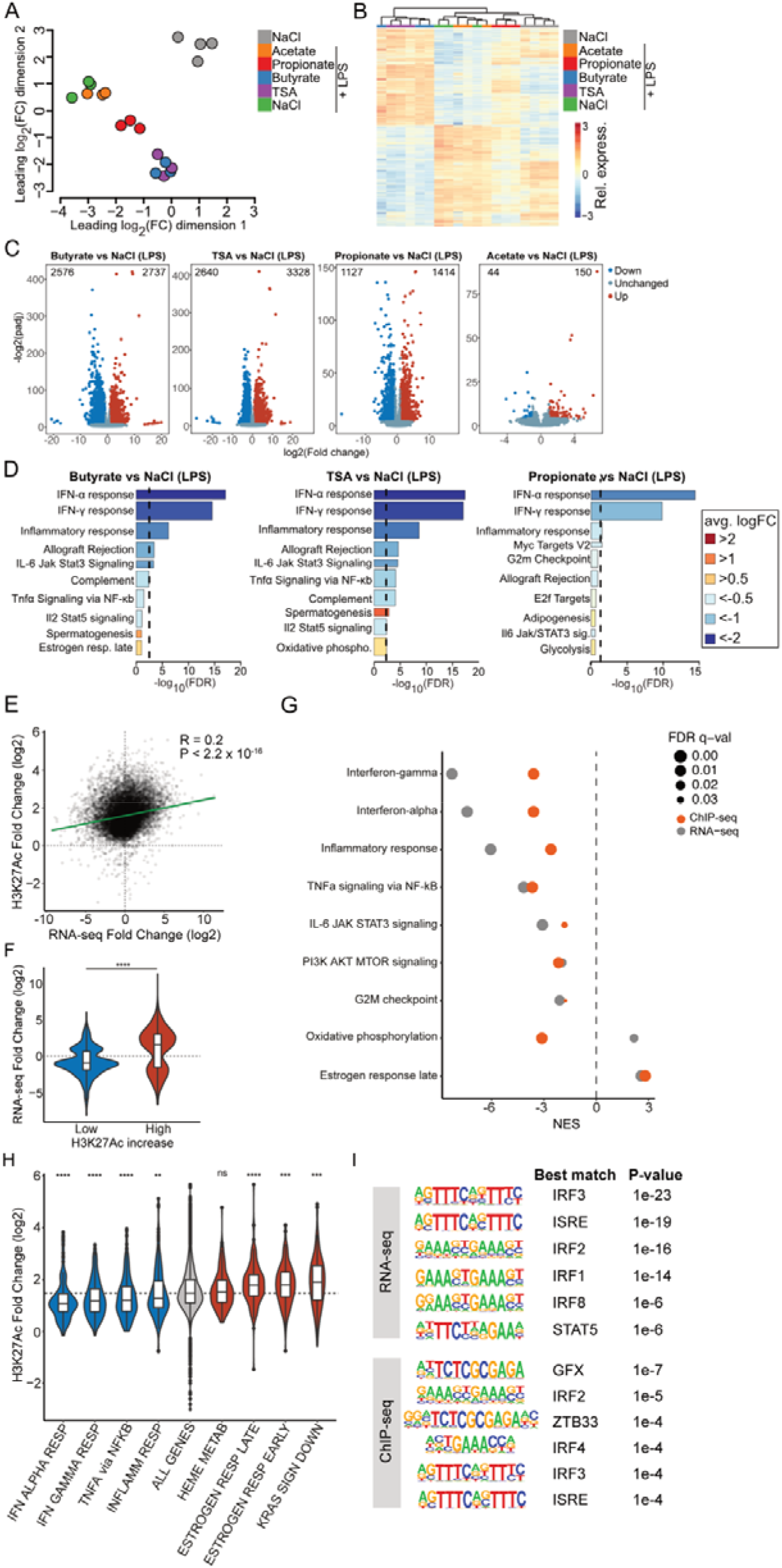
SCFAs alter the LPS-mediated transcriptome through HDAC inhibition, leading to histone hyperacetylation. RNA-seq analysis of hMDM were treated with either NaCl, acetate, propionate or butyrate (all 10 mM) or TSA (0.5 μM) in the presence of LPS (1 ng/ml) or left untreated for 16 h. (A) Multidimensional scaling (MDS) plot visualising the relationship between the samples, n = 3 or 4. (B) Heat map depicting relative expression values of transcripts that were significantly changed in LPS + butyrate vs LPS + NaCl comparison, scaled by row. (C) Volcano plots of genes altered by incubation with SCFAs or TSA compared to LPS alone. Significant up- and down-regulated genes (adjusted p-value < 0.05 and absolute log2(fold change) >1) are shown in red and blue, respectively. (D) Gene set enrichment analysis (GSEA) was performed based on the Molecular Signatures Database (MSigDB), using the hallmark gene sets. Plots show the top 10 hallmark gene sets. Bars are coloured by average log_2_(FC). Bar width represents the number of genes in the respective gene set. Dashed line indicates adjusted *p*-value threshold. (E) Correlation between the fold-change of differential H3K27ac signal at promoters (ChIP-seq) and the fold-change of differential gene expression (RNA-seq) comparing ‘Butyrate+LPS’ vs ‘LPS alone’ samples. Correlation was tested by Spearman correlation coefficient (R). (F) Distribution of the RNA-seq fold-change between ‘Butyrate+LPS’ vs ‘LPS alone’ for genes with the 10% lowest and 10% highest increase in H3K27ac ChIP-seq signal at promoters. (G) Normalized enrichment score (NES) from GSEA of hallmark gene sets using either RNA-seq or H3K27ac ChIP-seq signal for the comparison ‘LPS+Butyrate’ vs ‘LPS alone’. Only gene sets that were significant in both datasets (FDR q-val<0.05) are shown. (H) Fold-change in H3K27ac signal at promoters of genes belonging to gene sets negatively and positively enriched (by GSEA) in blue and red, respectively. In grey, values for all genes are shown. (I) Motif enrichment analyses for downregulated genes (RNA-seq) and the 10% of promoters with lowest H3K27ac increase (ChIP-seq) comparing ‘Butyrate+LPS’ vs ‘LPS alone’. *p<0.05, **p < 0.01, ***p < 0.001, ****p < 0.0001 by Wilcoxon’s test.

These findings suggested SCFA-mediated HDAC inhibition may alter transcription of inflammatory and interferon-related genes. Given that butyrate and propionate are known HDAC inhibitors, we investigated how butyrate altered histone acetylation. We first determined that butyrate increased total levels of acetylated lysine 27 on histone H3 (H3K27ac), a hallmark of active transcription at enhancers and promoters (Fig. S2C). We then delineated the genomic distribution of the increase in H3K27ac by performing chromatin immunoprecipitation sequencing (ChIP-seq), which was normalized with spiked-in chromatin to account for the global increase in acetylation. Consistent with the hyperacetylation observed by immunoblot, butyrate increased the number of the total called peaks by almost 4-fold (Fig. S2D), and caused a general increase in H3K27ac both at promoters and at intergenic peaks (Fig. S2E), consistent with the hyperacetylation observed by WB. However, the increase in H3K27ac at gene promoters was unevenly distributed and significantly correlated with the changes in gene expression caused by butyrate (Fig. 3E). Genes with the lowest increase in acetylation tended to show decreased transcription, and vice versa (Fig. 3F). Consistently, gene set enrichment analysis (GSEA) showed that pathways enriched among downregulated genes were also significantly enriched among genes with low acetylation increase, and included pathways such as ‘Interferon’, ‘Inflammatory response’ or ‘NF-kB’ (Fig. 3G). Genes belonging to these pathways had less acetylation increase than the average of all genes or specific subsets with high hyperacetylation (Fig. 3H). Transcription factor (TF) binding motif enrichment of downregulated gene promoters in RNA-seq (But + LPS vs. LPS) and the 10% of genes with lowest H3K27ac increase (But + LPS vs. LPS) showed a significant enrichment for interferon associated transcription factors (Fig. 3I). In contrast, enrichment for TF binding motifs from upregulated genes or those with the most increased H3K27ac showed general promoter-binding TFs and no consistent pathways (Fig. S2F).

To determine if the changes seen on the RNA level carried over to changes in protein levels, we performed proteomics on hMDM incubated with LPS +/- butyrate or TSA. While we identified considerably fewer differentially regulated proteins compared to the RNA-seq results (Fig. S2G), butyrate and TSA still clustered together by multidimensional scaling plot (Fig. S2H) and hierarchical clustering (Fig. S2I). GSEA showed that similar pathways to the RNA-seq were altered, demonstrating that the changes observed in the RNA-seq data were also occurring on protein level (Fig. S2J).

### SCFAs decrease expression of cFLIP and XIAP

The butyrate- and TSA-dependent decrease in genes involved in the inflammatory response was particularly interesting. Previous studies have shown that inhibition of the NF-kB signalling pathway, following engagement of a TLR, results in caspase-8 activation^27,28^, which can, in some cases, lead to activation of the NLRP3 inflammasome^29^.

We analysed the RNA-seq data set for changes in expression of proteins known to regulate caspase-8 activation and found that, in the context of LPS, butyrate and TSA decreased expression of two key regulators of caspase-8, cFLIP (*CFLAR*) and XIAP (Fig. 4A). Consistent with the global correlation between gene expression and H3K27ac changes induced by butyrate, both the *CFLAR* and *XIAP* genes were among the 10% with the lowest increase in promoter acetylation (Fig. 4B). We confirmed the decrease in expression by qPCR, which showed that butyrate and TSA decreased transcripts of both genes regardless of LPS pre-treatment (Fig. 4C).

**Figure 4.**
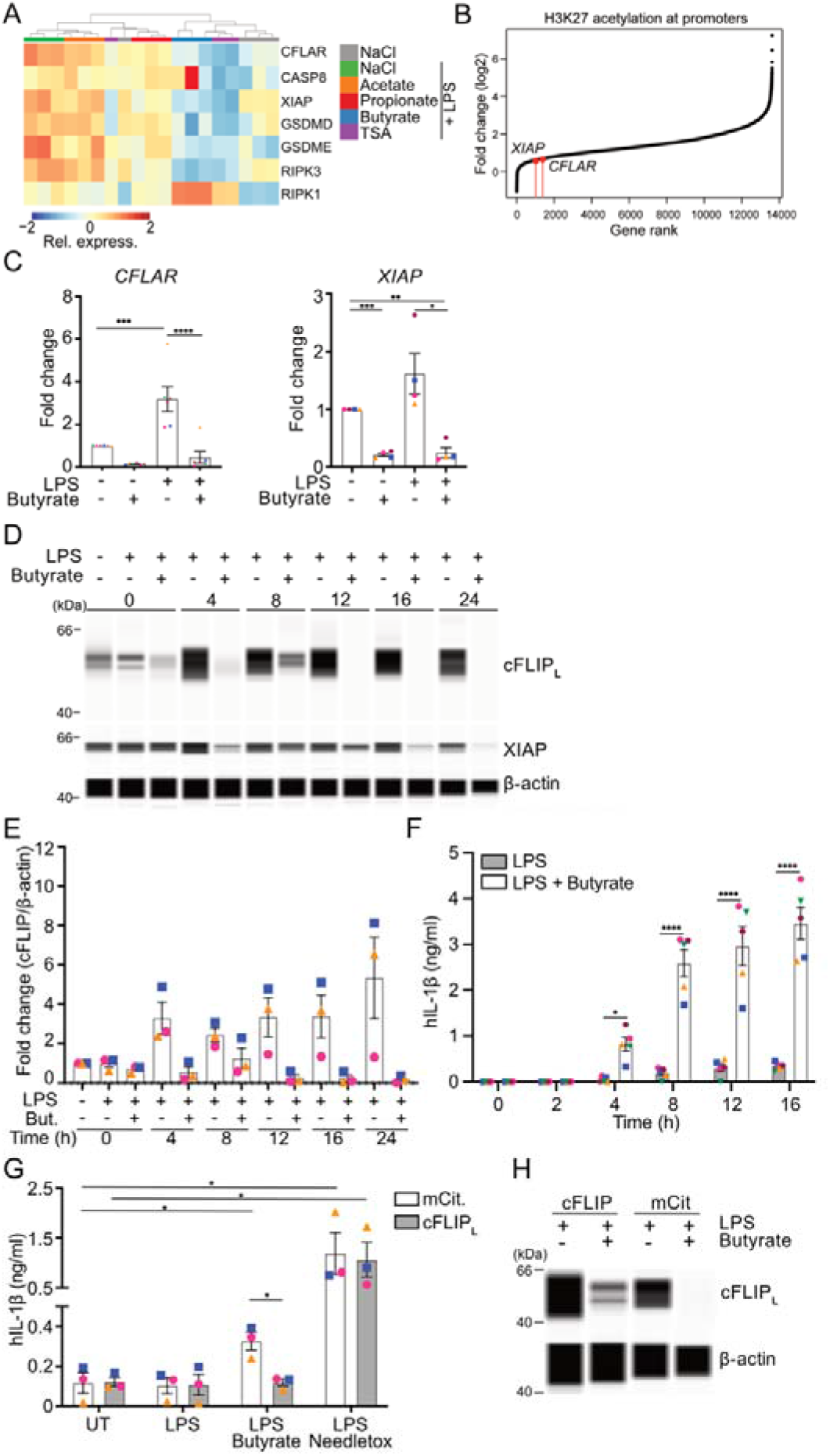
Butyrate-mediated decrease in cFLIP expression is required for NLRP3 inflammasome activation. (A) Heat map depicting expression of the relative subset of cell death related genes expression values of transcripts that were significantly changed in LPS + butyrate vs LPS + NaCl comparison, scaled by row (n=3). (B) Ranking of all genes ordered by the fold-change in H3K27ac signal in LPS + butyrate-treated samples. (C) *CFLAR* and *XIAP* expression assessed by qPCR. hMDM treated with LPS (1 ng/ml), or LPS and butyrate (10 mM) for the listed times and (C) cFLIP_L_ or XIAP assessed by WES (D) and quantitated in (E). Representative of 6 (B and C) independent experiments, and the blots shown (D) are representative of 4 independent experiments. (F) IL-1β release measured from cell-free supernatants from (D). (G) IL-1β release measure from hMDM transduced with cFLIP_L_ or vector alone and stimulated for LPS (1 ng/ml) and/or butyrate (5 mM, 16 h) or needletox (1 μg/ml needletox, 1 μg/ml PA, 1.5 h). (H) Immunoblot of hMDM from (G), representative of 3 independent experiments. Data are represented as mean +/- SEM, unless otherwise indicated, each dot is representative of one donor in all cases, *p<0.05, **p < 0.01, ***p < 0.001, ****p < 0.0001 by either one way ANOVA with Turkey (C) or Two-way ANOVA with SLJídák’s multiple comparisons test (F).

cFLIP can be transcribed in multiple isoforms, a long isoform (cFLIP_L_) or two short isoforms (cFLIP_R_ or cFLIP_S_)^30^. We assessed whether cFLIP was reduced upon LPS and/or Butyrate treatment by immunoblot and found that, consistent with alterations on the RNA level, expression of both the long and short isoforms of cFLIP were reduced in the presence of butyrate (Fig. S3A). We then correlated cFLIP and XIAP expression with IL-1β release over time and found that cFLIP_L_ was almost completely absent as early as 4 h, followed by a transient increase at 8 h, and then complete loss afterward (Fig. 4D, 4E). The loss of cFLIP correlated with the time we observed initial IL-1β release (Fig. 4F). In contrast, XIAP decreased across the time course but was not completely absent until 12-16 h (Fig. 4D, 4E). To ensure that loss of cFLIP was due to changes in gene expression, we assessed cFLIP degradation. However, inhibitors of the proteasome had no effect on cFLIP protein level in this time frame (Fig. S3B).

### Butyrate-dependent ablation of cFLIP expression is required to activate the NLRP3 inflammasome

The correlation between IL-1β release and the loss of cFLIP expression suggested that cFLIP would be regulating NLRP3 activation. To determine if loss of cFLIP was causal in SCFA-mediated NLRP3 activation, we transduced hMDM with lentivirus encoding either cFLIP_L_-T2A-mCitrine or mCitrine alone and stimulated them with butyrate and LPS. To ensure that we had close to 100% transduction efficiency we reduced the number of cells used per well. We determined that exogenous expression of cFLIP prevented LPS and butyrate-mediated IL-1β release (Fig. 4G), demonstrating that cFLIP was sufficient to inhibit NLRP3 activation. cFLIP was expressed at a comparable level to the endogenous expression in the absence of butyrate (Fig. 4H), confirming that this effect was not due to over-expression.

### Butyrate-mediated NLRP3 activation is dependent on caspase-8

The requirement for loss of cFLIP expression suggested that caspase-8 activation would be required for NLRP3 activation, and HDAC inhibition was demonstrated to trigger caspase-8 activation previously ^26^. Therefore, we assessed caspase-8 activity in hMDM following stimulation with LPS, butyrate or the combination and found that butyrate increased caspase-8 activity in both the absence or presence of LPS (Fig. 5A). To determine if it was required for NLRP3 activation, we pre-incubated hMDM with z-IETD-fmk, a capase-8 inhibitor, prior to stimulation with LPS and butyrate and determined that caspase-8 inhibition indeed significantly reduced NLRP3-dependent IL-1β release (Fig. 5B) and cleavage (Fig. 5C), as well as caspase-1 cleavage (Fig. 5D). We confirmed this result using siRNA against caspase-8, which partially ablated caspase-8 expression (Fig. 5E). Partial ablation of caspase-8 similarly reduced butyrate-mediated NLRP3 activation (Fig. 5F). Notably, VX-765 at the concentration used in our experiments did not inhibit caspase-8 activity, confirming that caspase-8 is upstream of NLRP3-inflammasome activation (Fig. S4A).

**Figure 5.**
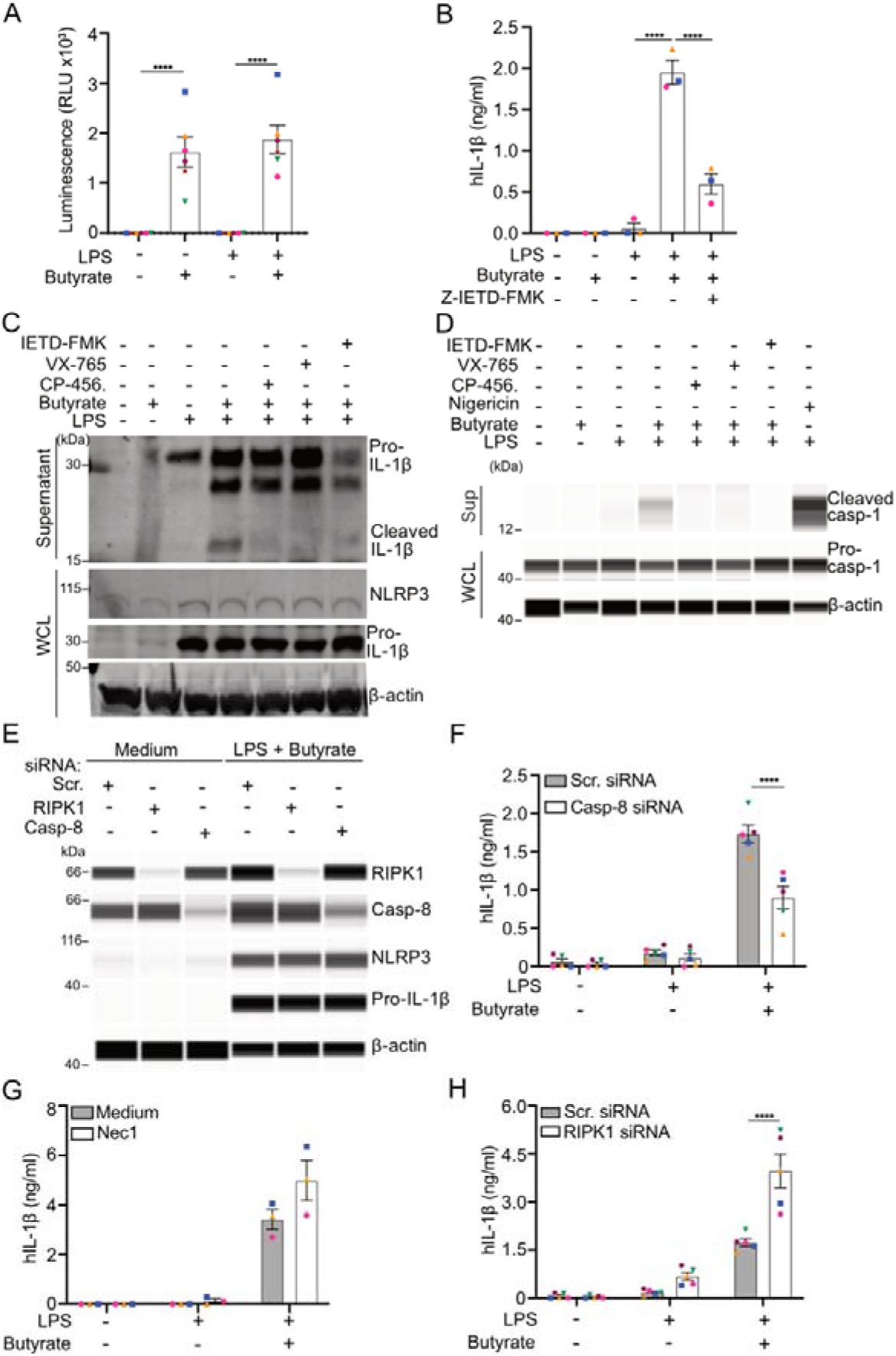
Butyrate-mediated NLRP3-inflammasome activation is dependent on caspase-8 and restrained by RIPK1. (A) hMDM were treated with medium, butyrate (10 mM) or LPS (1 ng/ml), alone or in combination, for 16 h and assessed for (A) The activity of the intracellular caspase-8 was assessed using Caspase-Glo^®^ assay kit, luminescence is proportional to caspase-8 activity. Data shown represents 6 independent experiments. (B) IL-1β in the cell-free supernatants from hMDM were treated with medium, LPS (1 ng/ml) or butyrate (10 mM), alone or in combination, for 16 h in the presence or absence of Z-IETD-FMK (5 μΜ) 30 min before treatment with activators. Represents 3 independent experiments. (C) Immunoblot of hMDM treated as in (B) in the presence or absence of CP- 456,773 (2 μΜ), VX-765 (40 μΜ) or Z-IETD-FMK (5 μΜ) 0.5 hour before treatment with activators. Sup is precipitated supernatant, whole cell lysates (WCL). Representative of 2 independent experiments. (D) Immunoblot of hMDM treated as in (C) blotted for caspase-1. Sup is precipitated supernatant, whole cell lysates (WCL). Representative of 2 independent experiments. (E) Immunoblot of WCL from hMDM electroporated with either scrambled siRNA, RIPK1 siRNA or caspase-8 siRNA before the stimulation with medium, LPS (1 ng/ml) or LPS (1 ng/ml) + butyrate (10 mM) for 16 h. Representative of 3 independent experiments. (F) IL-1β from cell-free supernatants from hMDM electroporated with either scrambled siRNA or caspase-8 siRNA then stimulated with medium, LPS (1 ng/ml) or LPS (1 ng/ml) + butyrate (10 mM) for 16 h. IL-1β release from cell-free supernatants from hMDM were treated with medium, LPS (1 ng/ml) or butyrate (10 mM), alone or in combination, for 16 h in the presence or absence of Necrostatin-1 (1 μΜ). (G) or following electroporation with siRNA against a scrambled control (scr. ctl) or RIPK1 (H). Data shown represent 5 (F and H) or 3 (G) independent experiments. Data are represented as mean +/- SEM, unless otherwise indicated, each dot is representative of one donor in all cases, *p<0.05, **p < 0.01, ***p < 0.001, ****p < 0.0001 by either one way ANOVA with Turkey (A and B) or Two-way ANOVA with SLJídák’s multiple comparisons test (F, G and H).

### Butyrate-mediated NLRP3 activation is independent of RIPK3 but restrained by RIPK1

Caspase-8 is activated by oligomerisation, which is facilitated by binding to the scaffolding adaptor protein FADD, upon engagement by the death domains of either activated RIPK1 or TRADD. We first investigated whether RIPK1 was required using both a RIPK1 inhibitor, Necrostatin-1 (Nec1) or using siRNA knock down, as RIPK1 has both kinase dependent and independent functions ^31^. Inhibiting RIPK1 kinase activity with Nec1 caused a minor and non-significant increase in NLRP3 activation by LPS and butyrate (Fig. 5G), demonstrating that RIPK1 activity is not required for NLRP3 activation, and suggests that it is not required for butyrate-dependent capase-8 activation. To determine whether RIPK1 had an enzymatic independent function, we used siRNA to knock down RIPK1. The RIPK1 knock down effectively reduced RIPK1 expression (Fig. 5E). Strikingly, knocking down RIPK1 substantially increased LPS and butyrate dependent IL-1β release, and enabled IL-1β release in response to LPS alone (Fig. 5F). This effect was restricted to IL-1β, as TNF-α was unchanged by both RIPK1 inhibition or knock down (Fig. S4B). This result was surprising, and demonstrates that RIPK1 is a negative regulator of both butyrate and spontaneous NLRP3 activation in hMDM. We then investigated RIPK3 by siRNA knock down and determined that even though the target could be ablated (Fig. S4C) it had no effect on butyrate-mediated NLRP3 activation (Fig. S4D).

### Loss of IL-10 secretion is required for butyrate-driven NLRP3 inflammasome activation

Alongside the increase of IL-1β release, butyrate decreased secretion of other cytokines including IL-6, IL-10 and IL-12p40 (Fig. 1A). IL-10, in particular, is important in maintaining gut homeostasis, and this is partially through inhibition of NLRP3 inflammasome activation^32^. In accordance with our sequencing and the cytokine bead array data, we determined that IL-10 secretion was completely blocked by co-incubation of LPS with butyrate, even as low as 1 mM, a similar concentration required to trigger IL-1β release (Fig. 6A). This was also consistent across a range of TLR ligands (Fig. S5A). Similar to *CFLAR* and *XIAP*, the *IL-10* gene locus had little increase in acetylation compared to other expressed genes following butyrate and LPS (Fig. 6B). Therefore, we tested whether loss of IL-10 was also due to HDAC inhibition and determined that inhibition of class 1 HDACs (Fig. 6C) was sufficient to inhibit IL-10 expression and secretion. To determine if loss of IL-10 secretion was a requirement for NLRP3 activation, we treated the cells with LPS and butyrate in presence or absence of recombinant IL-10. Pre-incubation with IL-10 completely ablated activation of NLRP3 with LPS and butyrate (Fig. 6D), demonstrating that loss of IL-10 or other cytokines that activate the same signalling pathways is required for butyrate-mediated NLRP3 activation.

**Figure 6.**
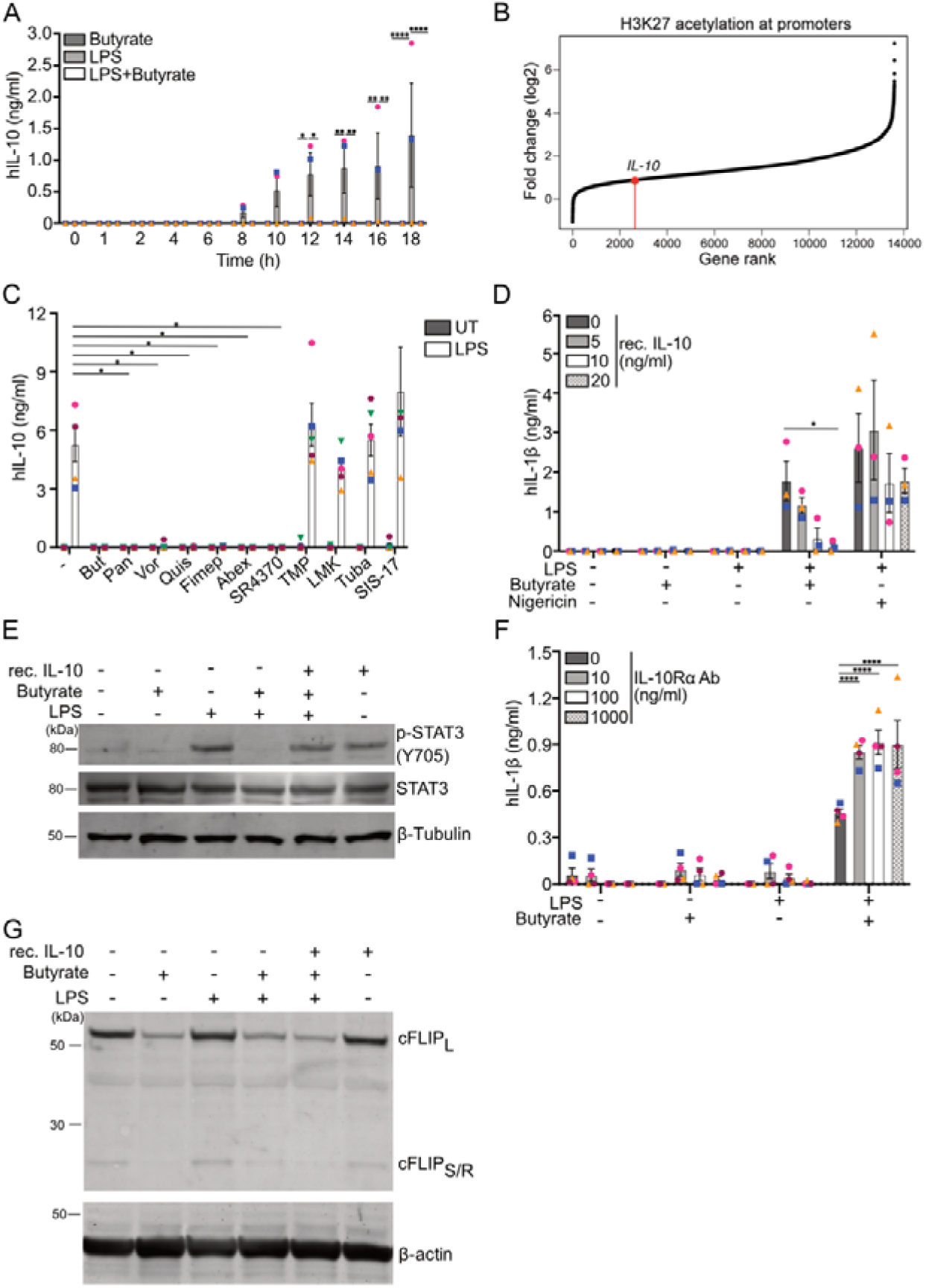
Loss of STAT3 activating cytokines is required for Butyrate-driven NLRP3 inflammasome activation. (A) IL-10 measured from cell-free supernatants from hMDM were treated with butyrate (10 mM) or LPS (1 ng/ml), alone or in combination, for the indicated period of time. Data shown represents 3 independent experiments. (B) Ranking of all genes ordered by the fold-change in H3K27ac signal in LPS + butyrate-treated samples. (C) IL-10 measured from cell-free supernatants from hMDM were treated with LPS +/- HDAC inhibitors. Data shown represents 5 independent experiments. (D) IL-1β release from hMDM incubated with LPS (1 ng/ml) and butyrate (10 mM) +/- pre-incubation with rhIL-10 (0.5 h). Data shown represents 3 independent experiments. (E) Immunoblot of phosphorylated STAT3(Y705), total STAT3 and β-Tubulin from hMDM pre-treated with rhIL-6 (100 ng/ml, 0.5 hour) or rhIL-10 (100 ng/ml, 0.5 hour) before subsequent treatments with medium, LPS (1 ng/ml) or butyrate (10 mM), alone or in combination, for 16 h. The blots shown are representative of 3 independent experiments. (F) IL-1β release from hMDM pre-incubated with IL-10Rα blocking antibody for 30 min, then LPS (1 ng/ml) or LPS (1 ng/ml) + butyrate (10 mM) for 16 h. Data shown represent 4 independent donors. (G) Immunoblot of phosphorylated cFLIP_L_, cFLIP_S_ and and β-actin from hMDM pre- treated with rhIL-10 (100 ng/ml, 0.5 hour) before subsequent treatments with medium, LPS (1 ng/ml) or butyrate (10 mM), alone or in combination, for 16 h. The blots shown are representative of 2 independent experiments. Data are represented as mean +/- SEM, unless otherwise indicated, each dot is representative of one donor in all cases, *p<0.05, **p < 0.01, ***p < 0.001, ****p < 0.0001 by Two-way ANOVA with SLJídák’s multiple comparisons test.

IL-10 was previously shown to inhibit NLRP3 through activation of STAT3 in a model of colitis ^21^. Therefore, we assessed whether STAT3, a known regulator of NLRP3 in the gut, was also inactivated by pre-treatment with butyrate and LPS. We determined that LPS resulted in phosphorylation of STAT3 at Tyr705, which was ablated by the addition of butyrate (Fig. 6E). This was also observed in response to activation of other TLRs (Fig. S6B). Notably, IL-10 rescued STAT3 phosphorylation at Tyr705 (Fig. 6E), demonstrating that butyrate inhibits STAT3 activation through inhibition of cytokine transcription.

To determine if loss of IL-10 signalling was sufficient to activate NLRP3, we pre-incubated hMDM with antibodies that blocked the IL-10 receptor (IL10Rα ab) or neutralised IL-10 (Neu. IL-10 ab) prior to stimulation. Inhibiting IL-10 signalling through either method increased IL-1β release in the presence of LPS and butyrate (Fig 6E, S5D), even at the lowest concentration used. However, crucially, neither enabled IL-1β release in the presence of LPS alone (Fig 6E, S5D). This was specific for IL-1β release, neither antibody altered TNFα secretion in either condition except at the highest concentration in the hMDM treated with LPS alone (Fig. S5C, D). We next determined whether STAT3 ablation would be sufficient to activate NLRP3 in hMDM through siRNA-mediated knock down. Both STAT3 targeting siRNAs effectively ablated STAT3 expression (Fig. S5E). However, whilst STAT3 knock down increased IL-1β release in the presence of LPS and butyrate but did not significantly enable IL-1β release by LPS alone, though some donors showed a small increase in IL-1β release (Fig. S5F). In contrast, TNFα secretion was unaltered by STAT3 knock down (Fig. S5F). These results collectively demonstrate that inhibiting IL-10 signalling or STAT3 signalling is insufficient to activate NLRP3 in this context.

In other contexts, IL-10 signalling increases expression of cFLIP through STAT3 activation ^33^, and so may be inhibiting butyrate-mediated NLRP3 activation by rescuing cFLIP expression. To determine whether this was the case, recombinant IL- 10 was added to hMDM together with LPS in absence or presence of butyrate and the cells were assessed for cFLIP expression. However, IL-10 did not increase expression of either form of cFLIP (Fig. 6F), demonstrating that its inhibition of butyrate-mediated NLRP3 activation is independent of cFLIP regulation.

### Butyrate-mediated NLRP3 activation does not trigger cell death but facilitates gasdermin D-independent IL-1**β** secretion

Having established the pathway used by butyrate to activate NLRP3, we next sought to determine the outcomes of butyrate-mediated NLRP3 activation for the cell. Our previous data had shown that while hMDM readily released IL-1β in response to LPS and butyrate they did not release LDH (Fig. 1C), suggesting that they do not undergo pyroptosis. To further clarify this, we assessed plasma membrane integrity using an exclusion dye in hMDM treated with LPS, butyrate or a combination of both over 16 h. LPS and nigericin was included as a positive control. Concordant with the lack of LDH release, hMDM treated with LPS and butyrate showed no dye uptake or changes in morphology (Fig. 7A), suggesting that they do not undergo cell death. This contrasted with the LPS and nigericin condition, where dye uptake occurred at an early-stage post treatment (Fig. 7A). This result was surprising, so we next assessed whether gasdermin D (gsdmD), the primary effector of pyroptosis, was cleaved in response to LPS and butyrate. However, we did not detect the active p30 C-terminal fragment of gsdmD, which was present in hMDM treated with LPS and nigericin. Instead, we detected the inactive p43 fragment of gsdmD (Fig. 7B) ^34^. To assessed the requirement for gsdmD functionally we used siRNA to knock it down (Fig. 7C), but this also had no effect on LPS and butyrate mediated IL-1β secretion (Fig. 7D), confirming its independence of gsdmD.

**Figure 7.**
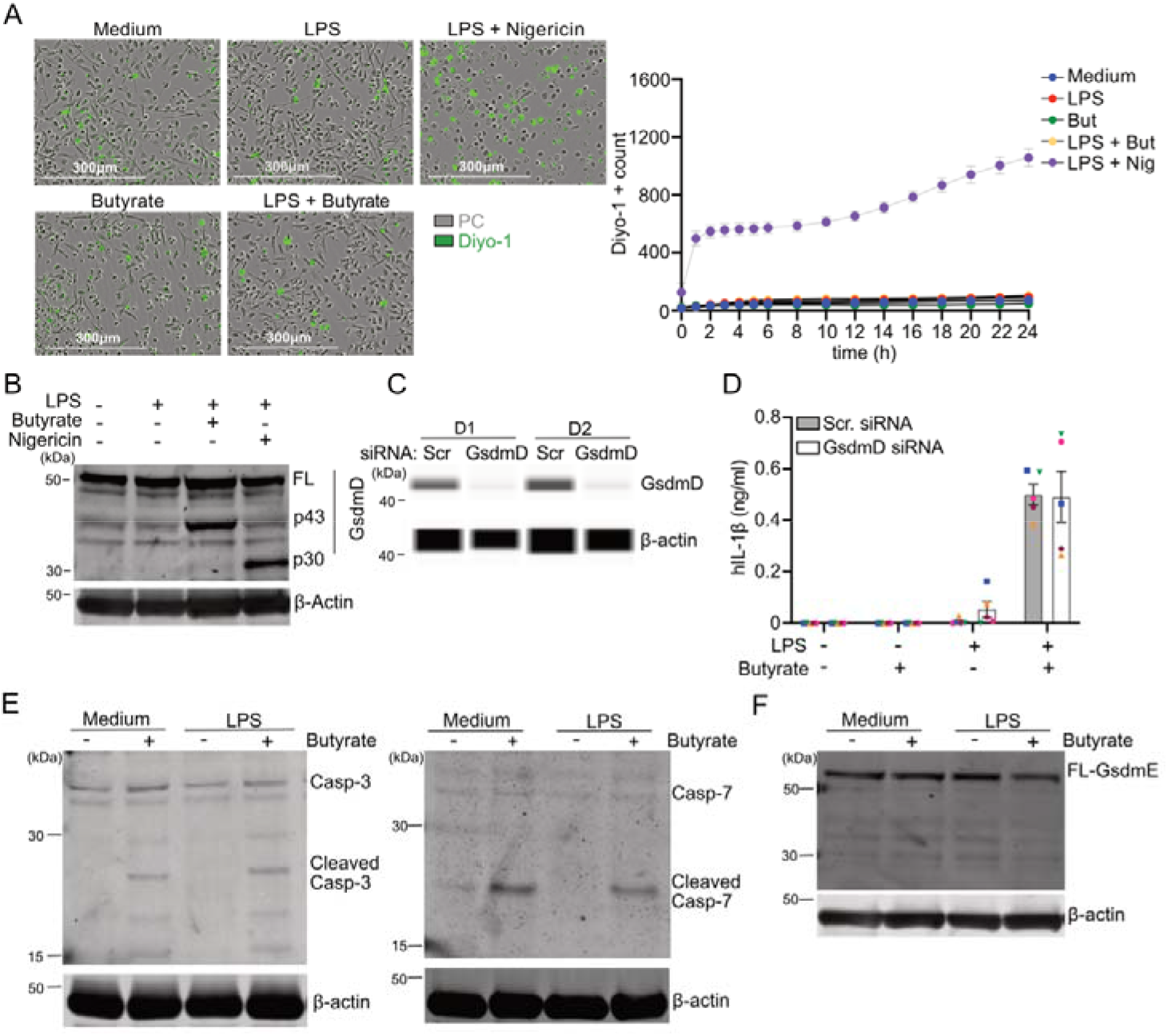
Butyrate-mediated NLRP3 activation triggers IL-1β secretion independently of gasdermin D and cell death. (A) hMDM were incubated with live cell imaging dye Diyo-1 (1:10,000) before treatment with medium, LPS (1 ng/ml) or butyrate (10 mM), alone or in combination, for 24 h, or with LPS (10 ng/ml, 3 h) and nigericin (10 μM). Cell death analysis was performed using the IncuCyte bioimaging platform, four images captured per well every one or two for 24 h. Representative images shown in upper panel of (A) taken at 24 h. Fluorescent dye Diyo-1 count/image in 24 h shown in the lower panel of (A) was averaged between four images/well. This was used to give average cell count/image/well, demonstrating membrane-permeabilized cells. Shown is one representative experiment out of n = 4 with technical duplicates. (B) hMDM were treated medium, LPS (1 ng/ml) or LPS (1 ng/ml) + butyrate (10 mM) for 16 h, or primed with LPS (1 ng/ml, 3 h) prior to subsequent administration of the NLRP3 inflammasome activator Nigericin (10 μM, 1.5 h). The levels of full length, cleaved form of gsdmD and β-actin, serving as loading control, were evaluated by Western blot. Blots are representative of three independent experiments. (C) Immunoblot from hMDM electroporated with either scrambled siRNA or gsdmD siRNA then stimulated with medium, LPS (1 ng/ml) or LPS (1 ng/ml) + butyrate (10 mM) for 16 h. (D) IL-1β from cell-free supernatants from hMDM treated as in (C). hMDM treated as in (B) but instead analysed for cleaved caspase-3 or cleaved caspase-7 (E) or gsdmE (F) by immunoblot. In all cases mean and SEM shown for 3-5 donors, each dot represents one donor. Immunoblots representative of 2 independent experiments.

The p43 fragment of gsdmD is formed when it is cleaved by caspase-3, which inactivates gsdmD. Therefore, we assessed caspase-3 and caspase-7 cleavage following incubation with LPS and butyrate and found the active, cleaved forms of both were present in the hMDM (Fig. 7E). Notably, activation of caspase-3 and -7 were independent of pre-incubation with LPS, as was observed for butyrate-mediated caspase-8 activation, demonstrating that multiple caspases are activated by butyrate. Gasdermins other than gasdermin D have been implicated in both cell death and release of IL-1β, including gasdermin E (gsdmE), which is cleaved and activated by caspase-3 ^34^. As caspase-3 was also active in these cells we assessed whether gsdmE was cleaved into its active form but found no difference in cleavage between the LPS condition and either butyrate alone or LPS and butyrate (Fig. 7E), indicating that gsdmE is also not required for IL-1β release.

## Discussion

SCFAs contribute to both the maintenance of the epithelial barrier and the programming of immune cells under steady-state conditions^35,36^. However, it is unclear what role SCFAs might play in the immune response under conditions where the intestinal barrier has become permeable. Our findings demonstrate that the effects of SCFAs depend on the context in which they are sensed. In contrast to the antimicrobial program engendered by SCFAs on macrophages in steady-state conditions, we show that butyrate and propionate instigate a pro-inflammatory program in the presence of pathogen- and danger-associated molecular patterns. This is primarily mediated through activation of the NLRP3 inflammasome and the subsequent release of IL-1β. NLRP3 could be activated by exposure to butyrate or propionate concurrent with stimulation of any TLR, demonstrating that translocation of any pathogen-derived molecules would be sufficient to trigger IL-1β release in their presence. Concurrently, butyrate and propionate reduced expression and secretion of many LPS-induced cytokines, including the anti-inflammatory molecules IL-10 and IL-1RA, amplifying the effect of NLRP3-dependent IL-1β release. Importantly, butyrate and propionate are produced exclusively by viable bacteria and are sensed by macrophages as danger signals. Therefore, under inflammatory conditions butyrate and propionate represent vita-PAMPs, microbial-derived products that signal bacterial viability ^37^. The requirement for ongoing synthesis of SCFAs is consistent with the two other vita-PAMPS characterised so far, cyclic-di- adenosine-monophosphate (c-di-AMP), and prokaryotic messenger RNA (mRNA), both of which need to be actively synthesised by live bacteria ^38^. The action of both butyrate and propionate as danger signals is further evidenced by the inhibition of a key IL-1 regulatory mechanism, IL-1RA, thus ensuring that the IL-1β-mediated pro-inflammatory response is propagated to trigger inflammation.

The vita-PAMP activity of butyrate and propionate is likely to be mediated by their ability to inhibit HDACs. Inhibition of HDACs, specifically HDACs 1-3 and 10, recapitulated both of the effects of butyrate and propionate, namely activation of NLRP3 and modulation of the LPS-mediated cytokine response. This differentiates them from the two previously described vita-PAMPs, which are detected directly by immune receptors. Notably, HDAC inhibition by either butyrate or TSA caused profound changes in the transcriptome, and in particular caused the loss of transcription of two important genes required to prevent NLRP3 inflammasome activation, *CFLAR* and *IL10*. Reconstitution or addition of either one of these molecules was sufficient to prevent NLRP3 inflammasome activation, demonstrating that butyrate and propionate engage a multifaceted program to enact NLRP3 inflammasome activation. It was interesting to note that, counterintuitively, loss of *CFLAR* and *IL10* expression occurred despite of increasing H3K27ac across the genome. This suggests that HDAC inhibition results in an altered pattern of histone acetylation caused by histone hyperacetylation, which may then overload the transcriptional machinery, resulting in it becoming the limiting factor for transcription. This has been observed previously for ISGs, where butyrate and other HDAC inhibitor cause a decrease in ISG expression^39^. This study demonstrated that histone hyperacetylation mediated by HDAC1/2 inhibition resulted in relocalisation of Brd4, an epigenetic reader, and the elongation factor P-TEFb, to other histones and away from ISG loci, causing loss of ISG expression. Given the correlation between the loss of expression of cFLIP and XIAP, and the lack of increased acetylation, we suggest that expression of these genes may be regulated by the same mechanism. The coupling of expression of ISG and interferon related genes, which are important for the response to pathogens, to genes essential or the prevention of cell death may be a mechanism to ensure the cell dies if expression of ISGs is interfered with. This is highly relevant to treatment for interferonopathies, where HDAC inhibitors have been proposed as a potential therapy. Further research exploring the link between alteration to histone acetylation and cell death could potentially lead to better understanding epigenetic modulation in both pathogen-mediated cell death and HDAC inhibitors as a therapy for auto-inflammatory and immune disorders.

In addition to inhibiting expression of cFLIP, butyrate-mediated NLRP3 activation also required a loss of IL-10 expression and secretion. This effect was also likely to be mediated by HDAC inhibition and subsequent loss of transcriptional machinery at the IL-10 locus, as it was also not highly hyperacetylated following treatment with butyrate. IL-10 regulated NLRP3 activation by controlling activation of STAT3, as loss of STAT3 in hMDM resulted in spontaneous NLRP3 activation following incubation with LPS. It was interesting to note here that inhibition of NLRP3 by IL-10 was independent of regulation of cFLIP, implying that the IL-10/STAT3 axis regulates NLRP3 activation independently of caspase-8 activity. This suggests that the presence of IL-10, rather than regulating cell death, will dictate whether caspase-8 dependent cell death is inflammatory or not, based on whether it activates apoptosis, an immunologically silent form of death, or leads to continued IL-1β release. It also demonstrates that SCFA-driven NLRP3 activation requires multiple, different perturbations to the LPS-transcriptome.

The loss of cFLIP expression enabled activation of caspase-8, which was required to subsequently trigger the NLRP3 inflammasome. Caspase-8 activation is a crucial cellular defence mechanism to guard against pathogen-mediated inhibition of the NF-kB-mediated inflammatory response ^40^. Indeed, inhibition of TAK1, an essential kinase in the NF-kB signalling pathway, is sufficient to trigger caspase-8 dependent cell death through loss of cFLIP expression^41^. However, beyond inhibition of this signalling cascade it is unclear how other mechanisms that interfere with expression of inflammatory genes triggers caspase-8 activation. Here we show that epigenetic-mediated loss of cFLIP is a new pathway through which this can occur. Further research focusing on pathogen-mediated loss of transcription and translation generally, or specifically for cFLIP, can lead to cell death and promote immunity.

The absence of potassium efflux, ASC speck formation and cell death accompanying butyrate-mediated NLRP3 activation was surprising. This is reminiscent of the alternative pathway of NLRP3 activation triggered by LPS in monocytes ^24^ or NLRP3 hyperactivation in response to ox-PAPC ^42^, where the cells also release IL-1β without the accompanying cell death. Indeed, despite butyrate-mediated activation of caspase-1, -3 and -7, IL-1β release was independent of gsdmD and gsdmE, consistent with the cells not becoming permeable to the exclusion dye, suggesting that pore forming proteins are not involved in IL-1β release. It is unclear how IL-1β is released in response to butyrate, though one possibility is the non-classical secretion pathway, which are reliant on the autophagy system and has been shown to release IL-1β in other contexts ^43^. Alternatively, it was recently shown that cleaved IL-1β could be translocated to the golgi and secreted through the canonical secretion pathway ^44^. Further research will be required to delineate the exact mechanism, as well as how the hMDM survive in spite of having multiple active caspases.

Our results also go some way to explaining the opposing effects of dietary fibre and SCFAs in IBD, as studies have suggested both beneficial and detrimental effects to the gut health in patients with IBD and in mouse colitis models ^11^. They suggest that the timing of the SCFA generation will have an impact on whether they function as anti- or pro-inflammatory. SCFA that are generated once the intestinal epithelial barrier has been breached and TLR ligands and pro-inflammatory cytokines including TNFα are present would result in NLRP3 inflammasome activation, thus contributing to the pathogenesis of IBD rather than supressing it. This would be exacerbated by the SCFA-driven loss of other cytokines, particularly IL-10, and the subsequent absence of STAT3 activation. Further research examining the role of fibre in different stages of IBD will help to elucidate in what contexts they are beneficial and when they may be detrimental.

## Material and Methods

### Experimental model and subject details

Buffy coats from healthy donors were obtained according to protocols accepted by the institutional review board at the University of Bonn (local ethics votes Lfd. Nr. 075/14).

### Isolation and culturing of primary human macrophages

Human PBMCs were obtained from buffy coat by density gradient centrifugation in Ficoll–Paque PLUS (GE Healthcare). PBMCs were incubated at 4 with magnetic microbeads conjugated to monoclonal anti-human CD14 Abs (Miltenyi Biotec) and isolated by positive magnetic selection according to manufacturer’s instructions (Miltenyi Biotec). Primary human macrophages were generated through differentiation of CD14^+^ monocytes with 500 U/mL of recombinant human GM-CSF (ImmunoTools) in RPMI 1640 medium (Life Technologies) containing 10% FBS (Thermo Fisher), 1% penicillin/streptomycin, 1 × GlutaMAX, and 1 × sodium pyruvate (both from Life Technologies) for 3 days. Human monocytes-derived macrophages (hMDM) were cultured in RPMI 1640 medium (Life Technologies) containing 10% FBS (Thermo Fisher), 1% penicillin/streptomycin, 1 × GlutaMAX, and 1 × sodium pyruvate.

### Inflammasome stimulation

For inflammasome stimulation assays hMDM were incubated with LPS (1 ng/ml) and SCFAs (0.1-10 mM) for 16 h prior to harvesting. For nigericin and needletox stimulation hMDM were stimulated O/N with LPS (1 ng/ml) before stimulation the following day with nigericin (8 uM) or needletox (Lfn-PrgI 10 ng/ml, PA 10 ng/ml) for 1.5 h prior to sample harvesting.

### Measurement of cytokine secretion

1 x 10^5^ hMDM/well were seeded in a 96-well plate in 100 μl. 50 μl of LPS and/or SCFAs (0.1-10 mM) were added simultaneously and the hMDM incubated for 16 h. 100 μl of cell-free supernatant was collected. The cytokine concentration was determined by HTRF or ELISA according to the manufacturer’s instructions. For some measurements, Bio-Plex Pro™ Human Cytokine 27-plex Assay kit was used according to the manufacturer’s instructions.

### Caspase-8 activity assay

1.5 x 10^5^ hMDM/well were seeded into white 96-well plate in 100 μl. 100 μl of the indicated stimuli were added to the cells. Cells were incubated for 16 hours. Afterwards, the plate was centrifuged for 5 minutes at 340 x g and supernatants were collected for the measurement of cytokines. The cells were washed once with PBS. Then 25 μl caspase-8 Glo assay buffer containing MG-312 were added to the cells immediately. After the samples were incubated for 30 – 60 minutes in dark, the luminescent signal was measured at 470 nm by a SpectraMax i3 system.

### Lactate dehydrogenase (LDH) assay

To determine the cell viability of hMDM in response to the treatment with LPS and butyrate, the activity of released LDH in the supernatants were measured and quantified as an indicator of cell death. 1 x 10^5^ hMDM were seeded into 96-well plates in 100 μl RPMI 1640 medium containing 0.1% FCS. Another 100 μl indicated stimuli were added to and cultured with the cells for 16 hours, after which the plate was centrifugated for 5 minutes at 340 x g to remove non-adherent cells from the supernatants. Meanwhile, 200 μl 1 x lysis buffer provided by the kit (Pierce LDH cytotoxicity Assay) were used to lyse the control cells as positive control. Afterwards, 25 μl supernatants and 25 μl LDH assay buffer were transferred and mixed in a transparent 384-well plate and incubate for 30 minutes in the dark. Then the absorbance was measured at 490 nm and 680 nm by a SpectraMax i3 system.

### Small interfering RNA (siRNA) electroporation in primary human macrophages

All the siRNA-mediated knockdown experiments in primary human macrophages were performed by electroporation using a Neon Transfection System (MPK5000; Invitrogen). Each reaction used 1.2 – 1.5 x 10^6^ hMDM, which were mixed with 10 μl buffer R and 75 pmol (1.5 μl) siRNA. Subsequently, the samples were taken up using a 10 μl Neon Pipette Tip and electroporated with the following protocol: 1400 V, 20 ms, 2 pulses. The electroporated cells were cultured in 10% FCS RPMI medium without antibiotics for 3 days. For some of the siRNAs with low knockdown efficiency, cells need to be electroporated again at day 3 after the first electroporation and cultured in 10% FCS RPMI medium without antibiotics for another 2 days. 0.5 – 1 x 10^6^ cells were lysed by 30 μl RIPA buffer containing cOmpleteTM EDTA-free protease inhibitor and phosphatase inhibitor to prepare the samples for the validation of siRNA-mediated knockdown efficiency by either Western blot or WES.

### Cloning

CFLAR WT was cloned from addgene plasmid #82936 by PCR before using restriction enzyme digest and ligation to insert it into a lentiviral vector containing a murine phosphoglycerate kinase 1 (PGK) promoter, as well as a C-terminal T2A followed by a mCherry and a Puromycin selection cassette.

### Lentiviral transduction of primary human macrophages

To produce the virus-containing supernatant 7x10^6^ HEK293T cells were plated in complete DMEM in a 10 cm dish. After 16 - 24 h, HEK293T cells were transfected with either a lentiviral construct encoding the gene of interest (2 µg per well) or a vector encoding the Vpx-Vpr restriction factors ^45^ with the pPAX2 (1 µg well), and pMD2.G (100 ng/well) plasmids using PEI MAX (Polyscience, 24765-100). Cells were incubated at 37C, 5% CO2 for approximately 12 h, and then the media was exchanged with RPMI containing 30% HI-FBS, and cells were incubated for another 36 h. After 36 h, the viral supernatant was collected using a 10 mL Luer-lock syringe attached to a blunt 18G needle and then filtered using a 0.45 mm filter unit into a 50 mL falcon. LentiX-concentrator was added as per manufactures instructions and the supernatant incubated overnight at 4°C. The virus was collected by centrifugation (1500xg, 50min, 4°C) and resuspended in complete RPMI. The Vpx-Vpr virus was mixed with the virus containing the gene of interest at a ratio of 1:1, 8 µg/ml polybrene was then added and the virus diluted 1:7 in complete RPMI. hMDM were seeded for transduction the day prior to transduction in 96 well plates, 0.4x10^5^ cells/well and adhered overnight. The virus containing media was added to the hMDM for approx. 14 hr at 37°C, 5% CO_2_. Following incubation, the cells were collected by centrifugation, and the virus-containing medium was removed and replaced by complete RPMI. The experiment was performed 24 h post-transduction.

### Cellular potassium quantification

Following treatment, the supernatant was removed from the hMDM, which were then washed 3x in potassium free balanced salt solution (125 mM NaCl, 5 mM N-methyl- glucamine chloride, 2 mM MgCl2, 1 mM CaCl2, 10 mM Glucose, 10 mM HEPES pH 7.4.) and lysed in 1 ml of ddH_2_O. The cells were freeze thawed three times to lyse them, then the supernatant harvested and clarified by centrifugation. Potassium was measured from the clarified supernatant by ICP-MS.

### Immunoblotting

Up to 1 × 10^6^ cells were lysed using RIPA buffer (20 mM Tris-HCl pH 7.4, 150 mM NaCl, 1 mM EDTA, 1% Triton X-100, 10% glycerol, 0.1% SDS and 0.5% deoxycholate) supplemented with PhosSTOP and complete protease inhibitors (Roche). Cell lysates were clarified by centrifugation at 10,000 x g for 10 mins, followed by the protein concentration quantification by BCA assay. Equal amounts of samples for immunoblotting in NuPAGE LDS sample buffer and reducing agent were always incubated at 85 °C for 10 min before loading onto a 4–12% or 10% NuPAGE Bis-Tris gel. As a size marker, 3 μl of PageRulerTM Plus prestained protein ladder were loaded on each gel. Samples were separated under denaturing and reducing electrophoretic conditions at 150 V in MOPS or MES buffer. Separated proteins were then transferred from the gel to an Immobilon-FL polyvinylidene difluoride (PVDF) membrane at 32 V for 1–1.5 h. Non-specific antibody binding to the membrane was blocked by incubation of the membrane with blocking buffer for at least 1 h at RT. Antibodies diluted in binding buffer were incubated with the blocked membrane overnight at 4°C. Membranes were washed three times with TBST and then incubated with the secondary antibody for 1 h at RT in binding buffer. Membranes were then washed twice in TBST and one last time in TBS before being developed on an Odyssey Scanner (LI-COR).

### Simple Western assay on WesTM

For some experiments, the whole cell lysates were resolved and analyzed using the Simple Western size assay and Wes module. Protein concentration of each sample was normalized using the BCA assay to 1–2 μg/μl. Four parts of the lysates were combined with one part of the 5x Fluorescent Master Mix containing a fluorescently labelled standard, DTT and Sample Buffer, and were denatured by incubation at 95 °C for 5 min. The samples, a biotinylated ladder, the primary antibody, streptavidin-horseradish peroxidase (HRP), the secondary antibody, the chemiluminescent substrate, and wash buffer were loaded, as indicated by the manufacturer, into a microplate pre-filled with Split Running Buffer, Wash Buffer and 10x Sample Buffer. Samples were electrophoretically separated and detected in a 25-capillary cartridge (12–230 kDa) using the following protocol: 200 s loading time of the separation matrix, 15 s loading time of the stacking matrix, 9 s loading time of the sample, 25 min separation time of the sample at 375 V, 90 min incubation with the primary antibody and 30 min incubation with the secondary antibody. Results were analyzed with the Compass for Simple Western software.

### Live cell imaging for cell death analysis

hMDM were used to determine cell viability by live cell imaging. 3.5 x 10^4^ cells /well were seeded into a 96-well plate in RPMI 1640 medium containing rhGM-CSF (3.1 μl/ml). Diyo-1 (1:10,000), a live cell imaging dye, was also added. Cells were allowed to reattach for 4 hours before addition of any compounds. Cell death analysis was performed after stimulating the cells with LPS (1 ng/ml) and butyrate (10 mM), alone or in combination, using the IncuCyte bioimaging platform, which is housed in a high humidity, 37°C, 5% CO_2_ incubator. LPS (10 ng/ml) was added 3 hours prior to addition of nigericin (10 µM) as a positive control. Four images were captured per well in the appropriate fluorescent channels and phase contrast every one or two hours for 24 hours. These images were analyzed using the IncuCyte analysis software. The fluorescent count/image was averaged between four images/well. This was used to yield the average cell count/image/well, demonstrating membrane-permeabilized cells in the case of Diyo-1. Each condition was measured in technical duplicates for each donor.

### ASC speck imaging and quantitation

200 μl of 1 x10^6^/ml hMDM per well were seeded in 8-well ibidi slides. The cells were treated with medium, LPS (1 ng/ml) or butyrate (10 mM), alone or in combination, for 16 hours, or primed with LPS (10 ng/ml, 3 h) prior to the addition of NLRP3 inflammasome activator Nigericin (10 μM, 1.5 h). Then, the cells were fixed with 200 μl 4% formaldehyde in PBS after 30 min incubation at room temperature, followed by two washes with 200 μl PBS. 100 μl of mixtures containing 10 μl human FcR blocking reagent and 90 μl permeabilization buffer were incubated with the cells for 10 minutes at 37°C. Afterwards, the cells were stained with 4 μl of directly labelled anti-ASC-647 (1:25) and incubated overnight at 4°C in the dark. To wash out the extra antibodies, the cells were washed twice with 200 μl permeabilization buffer. Subsequently, 100 μl DNA dye Hoechst diluted by PBS (1:3000) were incubated with the cells for 10 minutes at room temperature in the dark, which is followed by two washes with 200 μl PBS to remove the remaining Hoechst solution. Finally, the cells were filled with 200 μl PBS and imaged by Observer Z1 epifluorescence microscope (ZEISS). The analysis of ASC specks was processed by counting the number of cells using Hoechst as a nuclear marker as well as the ASC 647 signal to count the specks to ultimately calculate the number of ASC specks per cell. This analysis was performed by using the image analyser software Cell profiler 3.0.

### Gene expression analysis by quantitative PCR (qPCR)

1 x 10^6^ hMDM were seeded in 12-well plate and incubated with indicated stimuli for 16 hours, after which 350 μl RLT lysis buffer containing 1% (v/v) β-mercaptoethanol were used to lyse the cells at room temperature. RNA was isolated according to the manufacturer’s instructions (RNeasy Mini Kit, Qiagen). Equal quantities of RNA (500 -1000 ng) from each sample were reverse transcribed into cDNA using the SuperScript III Reverse Transcriptase and oligo dT_(18)_ primers. The qPCR was performed on the QuantStudio 6 Flex real time-PCR system, and the relative expression of the target mRNA was analysed using the ΔΔ CT method with HPRT as the reference mRNA. For primer sequences see Table S1.

### RNA-sequencing

1 x 10^6^ GM-MDMs were treated NaCl (10 mM), SCFAs (acetate, propionate or butyrate; 10 mM) or TSA (0.5 μM) in the presence or absence of LPS (1 ng/ml) for 16 hours and subsequently lysed in TRIZOL (Invitrogen) and total RNA was extracted using the RNeasy Mini Kit (Qiagen) according to the manufacturer’s protocol. RNA was eluted in RNase-free water. The quality of the RNA was assessed by measuring the ratio of absorbance at 260 nm and 280 nm using a Nanodrop 2000 Spectrometer (Thermo Scientific) and by visualization of 28S and 18S band integrity on a Tapestation 2200 (Agilent). Total RNA was converted into libraries of double-stranded cDNA molecules as a template for high-throughput sequencing using the Illumina TruSeq RNA Sample Preparation Kit v2. Briefly, mRNA was purified from 100 – 500 ng of total RNA using poly-T oligo-attached magnetic beads. Fragmentation was carried out using divalent cations under elevated temperature in Illumina proprietary fragmentation buffer. First strand cDNA was synthesized using random oligonucleotides and SuperScript II. Second strand cDNA synthesis was subsequently performed using DNA Polymerase I and RNase H. Remaining overhangs were converted into blunt ends via exonuclease/polymerase activities and enzymes were removed. After adenylation of 3′ends of DNA fragments, Illumina adaptor oligonucleotides were ligated to prepare for hybridization. DNA fragments with ligated adaptor molecules were selectively enriched using Illumina PCR primers in a 15 cycles PCR reaction. Size-selection and purification of cDNA fragments with preferentially 200 bp in insert length was performed using SPRIBeads (Beckman-Coulter). Size distribution of cDNA libraries was measured using the Agilent high sensitivity DNA assay on a Tapestation 2200 (Agilent). cDNA libraries were quantified using KAPA Library Quantification Kits (Kapa Biosystems). After cluster generation on a cBot, 75 bp single read sequencing was performed on a HiSeq1500 and de-multiplexed using CASAVA v1.8.2.

### RNA-sequencing analysis

Pre-processing of RNA-Seq data was performed by a standardized and reproducible pipeline based on the Docker system (Docker image is available via docker hub, limesbonn/hisat2). Briefly, alignment to the human reference genome hg19 from UCSC was conducted by Hisat2 (Hisat2, 2.0.6) (Kim et al., 2015) using standard settings. The external gene names and Entrez gene IDs matching the original Ensembl gene IDs were obtained using The biomaRt package (v2.38.0) (Kinsella et al., 2011) and A DGEList object was created using the raw counts and gene annotation using the edgeR package (v3.24.0) (Robinson et al., 2009). Since genes with very low counts are not useful, only genes that had at least 10 reads in a worthwhile number of samples determined by the design matrix were kept. In addition, each kept gene is required to have at least 15 reads across all the samples. Filtering was performed using the filterByExpr function. Afterwards, counts were normalised. The aim of normalisation is to remove systematic technical effects that occur in the data to ensure that technical bias has minimal impact on the results. Normalisation was done by using TMM method (Robinson and Oshlack, 2010) (weighted, trimmed mean of M-values) with the application of edgeR calcNormFactors function. In contrast to other procedures, where the proportion of each gene’s reads is computed relative to the total number of reads and compared across all samples, here it is taken into account that different experimental conditions might express a diverse RNA repertoire and therefore might lead to not directly comparable proportions. To put it simply, normalisation is supposed to level the median of gene expression values across samples, assuming that the majority of genes are expressed at an equal level. After data normalisation and with the edgeR package v3.24.0, estimateDisp was used to estimate common dispersion and tagwise dispersion and transform the data for linear modelling. Multidimensional scaling (MDS) plot visualising the relationship between the samples were displayed with the batch-corrected data according to all the donors. For each comparison, differentially expressed genes (DEGs) were identified with a false discovery rate (FDR)-adjusted p-value < 0.05. Differential expression analysis was also performed with DESeq (v. 1.34.0)^46^ to ensure robustness, and for comparison with ChIP-seq results. Gene set enrichment analysis was performed using the camera function of the limma package v3.38 (Wu and Smyth, 2012) with the hallmark gene set collections from the Molecular Signatures Database v5.2 (Liberzon et al., 2015; Subramanian et al., 2005).

### ChIP-seq

10 x 10^6^ GM-MDMs were treated with butyrate (10 mM) in the presence or absence of LPS (1 ng/ml) for 6 hours. They were cross-linked with 1% formaldehyde, lysed and sonicated in 1% Triton, 0.1% sodium deoxycholate, 0.5% SDS, 0.2 M NaCl, 10 mM Tris, pH 7.5, 10 mM EDTA, and 1X protease inhibitor cocktail (Roche 11836170001). Sonication was performed in a Covaris M220 (75% PIP, 200 CPB, 7°C, 10% DF) for 15 min in lysis buffer with 0.5% SDS. Lysates were incubated with 4 µg of H3K27ac antibody (Abcam ab4729) overnight at 4°C. For normalization, 50 ng of *Drosophila* spike-in chromatin (53083, Active Motif) and 2 µg of spike-in antibody (61686, Active Motif) were added. Antibody-bound chromatin was pulled down with protein G Dynabeads (Invitrogen 10003D). Samples were washed, RNase-treated and reverse cross-linked by incubation at 65°C in 1% SDS, 0.1 M NaHCO_3_ and proteinase K. DNA was purified using ChIP DNA Clean & Concentrator Kit (Zymo). Libraries were prepared using the NEBNext Ultra DNA Library Prep kit (E7645, New England Biolabs).

### ChIP-seq data processing

ChIP-seq libraries were sequenced along with input libraries as paired end 150bp reads. Reads were aligned to human genome hg38 and to fruit fly genome dm6 using Bowtie2 version 2.4.4 with ‘--very-sensitive’ flag set and a minimum fragment length (-X flag) of 1000bp^47^, all other parameters set to default. Normalization based on spike-in chromatin was performed within each biological replicate as follows: scaling factors were calculated for each sample as the ratio of total uniquely aligned counts to Drosophila relative to the counts in the sample containing the least number of Drosophila counts, and reads were down-sampled according to this ratio. Quality of the ChIP-Seq libraries were assessed using ChIPQC. Duplicate reads were identified using Picard MarkDuplicates and excluded from the downstream analysis. Genome wide coverage tracks in bigwig format were generated using DeepTools bamCoverage (v. 3.3.1)^48^. ChIP-Seq Peaks were identified by MACS2 (v. 2.2.5)^48^ using input libraries as control. Average ChIP-seq signal around TSS and intergenic peaks (>5 kb from closest gene) were calculated and plotted with deepTools using bigwig files as input (v. 3.3.1, computeMatrix and plotProfile, respectively). To quantify H3K27ac signal at promoters, ChIP-seq reads were quantified at promoters (+-500 bp of transcription start site) of all genes previously analyzed by RNA-seq. Differential ChIP-signal between samples was analysed with DESeq2 (v. 1.34.0)^46^, omitting default normalization as ChIP-seq signal was already normalized by spike-in chromatin.

### GSEA

GSEA was carried out using ranked gene lists based on RNA-seq expression (wald statistics from DESeq2 results) and on ChIP-seq signal (log2FC of ChIP-seq counts at gene promoters). GSEA software v. 4.2.2 was used with a ‘classic’ scoring scheme and MSigDB gene sets ^49^.

### Motif enrichment

Enrichment analyses of known transcription factor motifs were performed at promoters of up and downregulated genes in RNA-seq and at the 10% of promoters with highest and lowest H3K27ac increase. Homer’s findMotifsGenome.pl program ^50^ was utilized, using in each case all other promoters as background and a region size of 400 bp, with all other parameters set as default.

### Mass Spectrometry-based Proteomics (MS)

1.5 x 10^6^ GM-MDMs were cultured in 1 ml RPMI 1640 medium in 12-well plates and challenged by indicated stimuli for 16 hours. The cells were washed twice with PBS before lysing the cells with 200 μl SDS lysis buffer containing freshly added DTT. Cell lysates were collected into new Eppendorf tubes and boiled for 10 minutes. 800 μl pre-cooled acetone were mixed with the samples and then incubated overnight at -20°C. Subsequently, the samples were washed twice with 80% acetone after centrifugation (19,000 x g, 20 minutes, 4°C). The pellets at the bottom of tubes were dried for 15 minutes at room temperature and stored at -80°C.

Frozen samples were resuspended in 50□µl of digestion buffer containing 1% SDC, 10 mM TCEP, 55 mM CAA, 25 mM Tris (pH = 8) and boiled for 10□minutes to denature proteins. After sonication using a Bioruptor (Diagenode), protein concentration was measured via BCA assay. 50 ug of proteins were digested with 1 ug Lys-C and Trypsin overnight at 37°C and 1500 rpm. Peptides were desalted and purified using 2 discs of SDB-RPS material and re-suspended in 2% acetonitrile/0.1% TFA for LC-MS.

Reverse phase chromatographic separation of peptides was performed by loading approximately 200 – 500 ng of peptides on a 50-cm HPLC-column (75-μm inner diameter; in-house packed using ReproSil-Pur C18-AQ 1.9-µm silica beads; Dr Maisch GmbH, Germany) coupled to an EASYnLC 1200 ultra-high-pressure system Peptides were separated with a buffer system consisting of 0.1% formic acid (buffer A) and 80% acetonitrile in 0.1% formic acid (buffer B) using a linear gradient from 5 to 30% B in 110□minutes. The column temperature was set to 60°C.

The LC was coupled to a quadrupole Orbitrap mass spectrometer (Q Exactive HFX, Thermo Fisher Scientific, Rockford, IL, USA) via a nano-electrospray ion source. The mass spectrometer was operated in a data-dependent acquisition mode, collecting MS1 spectra (60,000 resolution, 300 –1650□m/z range) with an automatic gain control (AGC) target of 3E6 and a maximum ion injection time of 20□ms. The top-15 most intense ions from the MS1 scan were isolated with an isolation width of 1.4□m/z. Following higher-energy collisional dissociation (HCD) with a normalized collision energy (NCE) of 27%, MS2 spectra were collected (15,000 resolution) with an AGC target of 5E4 and a maximum ion injection time of 28□ms. Dynamic precursor exclusion was enabled with a duration of 30□s.

### MS data processing and analysis

Mass spectra were searched against the 2019 Uniprot mouse databases using MaxQuant version 1.5.5.2 with a 1% FDR at the peptide and protein level. Peptides required a minimum length of seven amino acids with carbamidomethylation as a fixed modification, and N-terminal acetylation and methionine oxidations as variable modifications. Enzyme specificity was set as C-terminal to arginine and lysine using trypsin as protease and a maximum of two missed cleavages were allowed in the database search. The maximum mass tolerance for precursor and fragment ions was 4.5 ppm and 20 ppm, respectively. ‘Match between runs’ was enabled to transfer peptide identifications between individual measurements with a 0.7-min window after retention time alignment. Label-free quantification was performed with the MaxLFQ algorithm using a minimum ratio count of 2. Protein identifications were filtered by removing matches to the reverse database, matches only identified by site, and common contaminants. Data filtering and Statistical analysis was performed with Perseus v1.5.5.5, GraphPad Prism v7.03, Microsoft Excel, and R Studio v3.4.0. Data was filtered further such that only proteins with identifications in all replicates of one cell type were retained. Missing values were imputed from a normal distribution of intensity values at the detection limit of the mass spectrometer. Statistical analysis was performed as indicated in the Figure legends with a constant permutation-based FDR correction at 5%.

### Statistical analysis and quantification

Data analysis and statistical analysis was performed in R and GraphPad Prism. All statistical analysis were preceded by Normality tests, followed by the recommended parametric or nonparametric tests. If not stated otherwise, P values were determined by two-way ANOVA with Tukey’s or Sidak’s or Bonferroni’s Values of p < 0.05 were considered statistically significant. *p < 0.05, **p < 0.01, ***p < 0.0002, ****p<0.0001. Data are graphed as mean; error bars show SEM unless otherwise stated.

## Key resources table

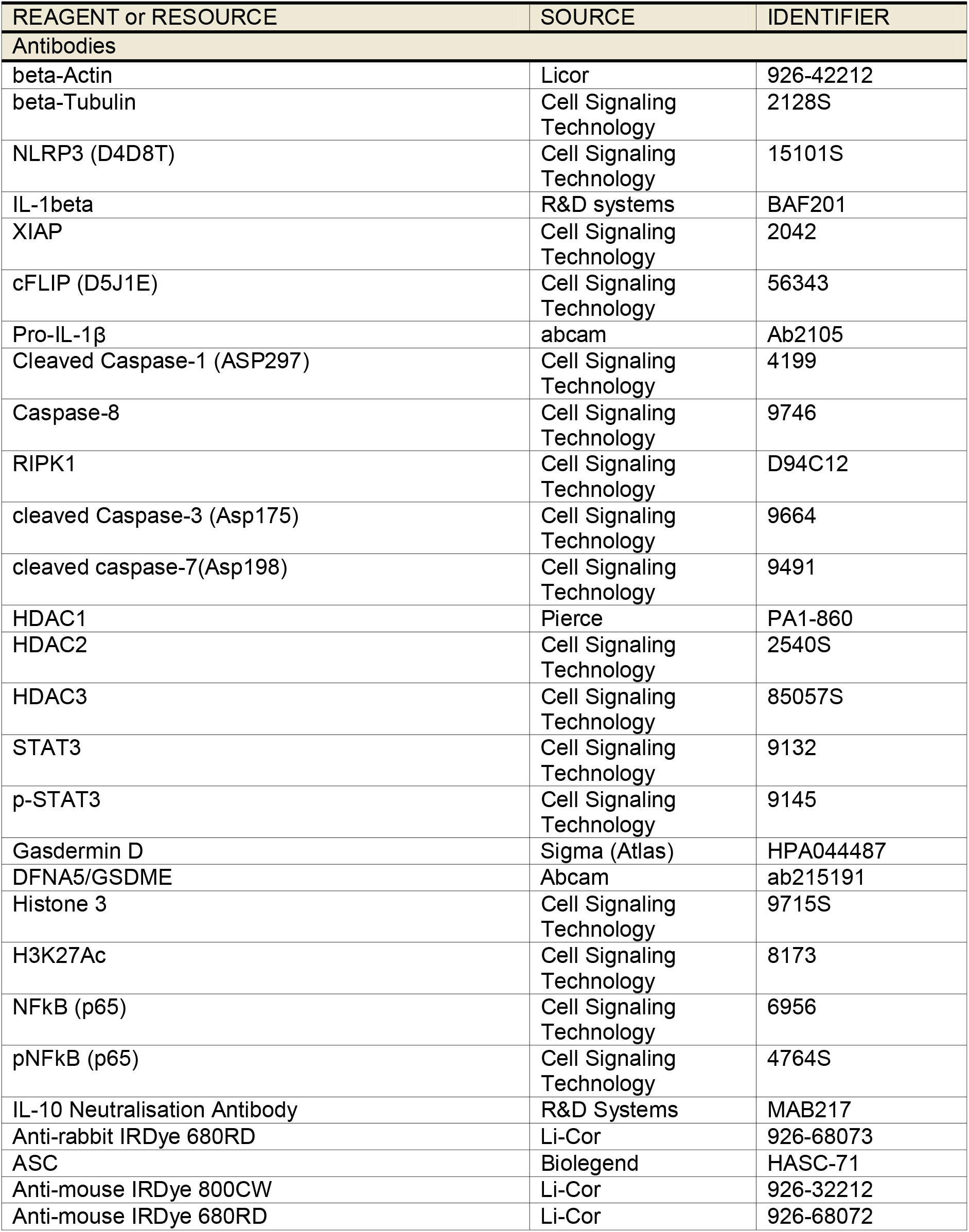
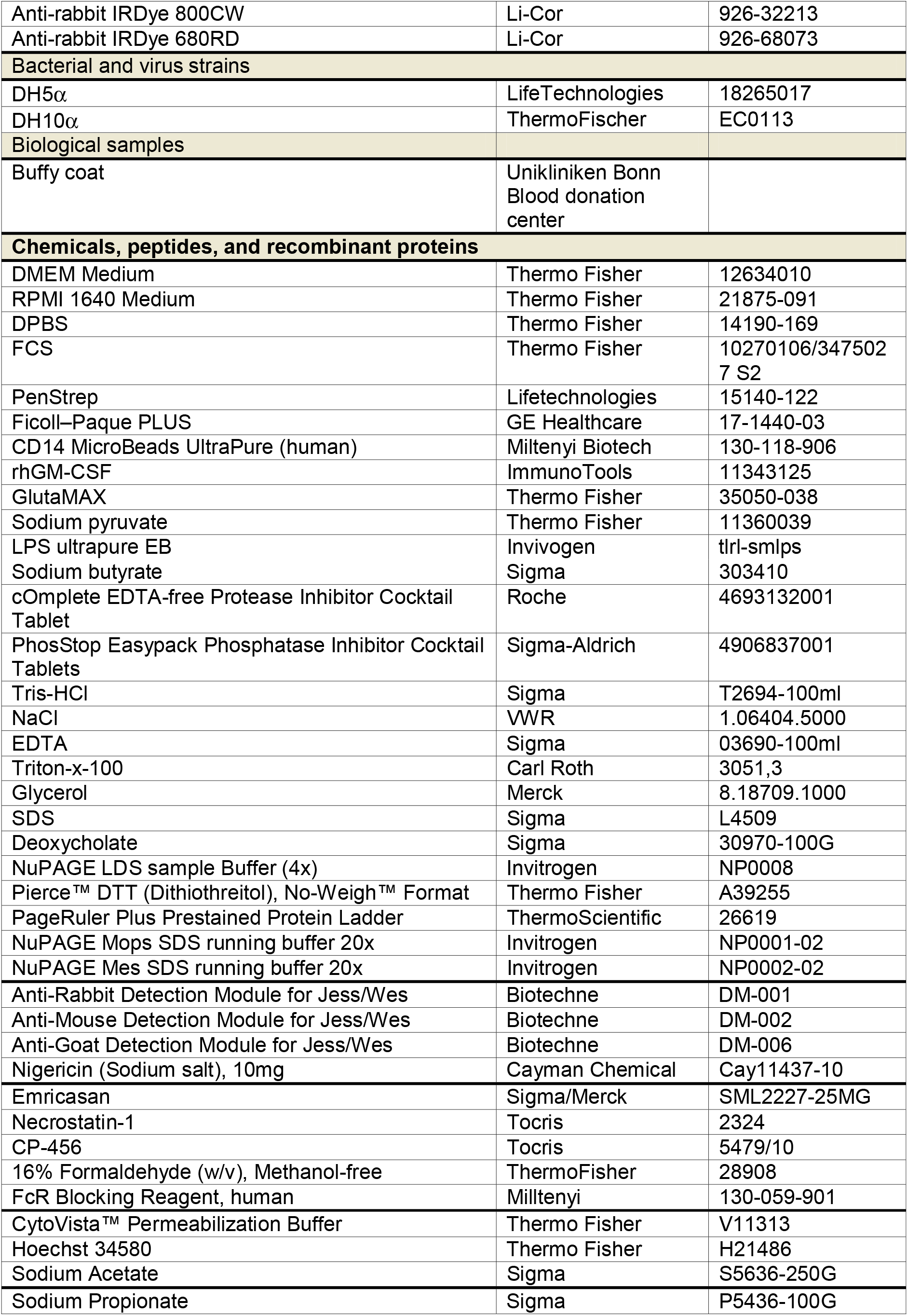
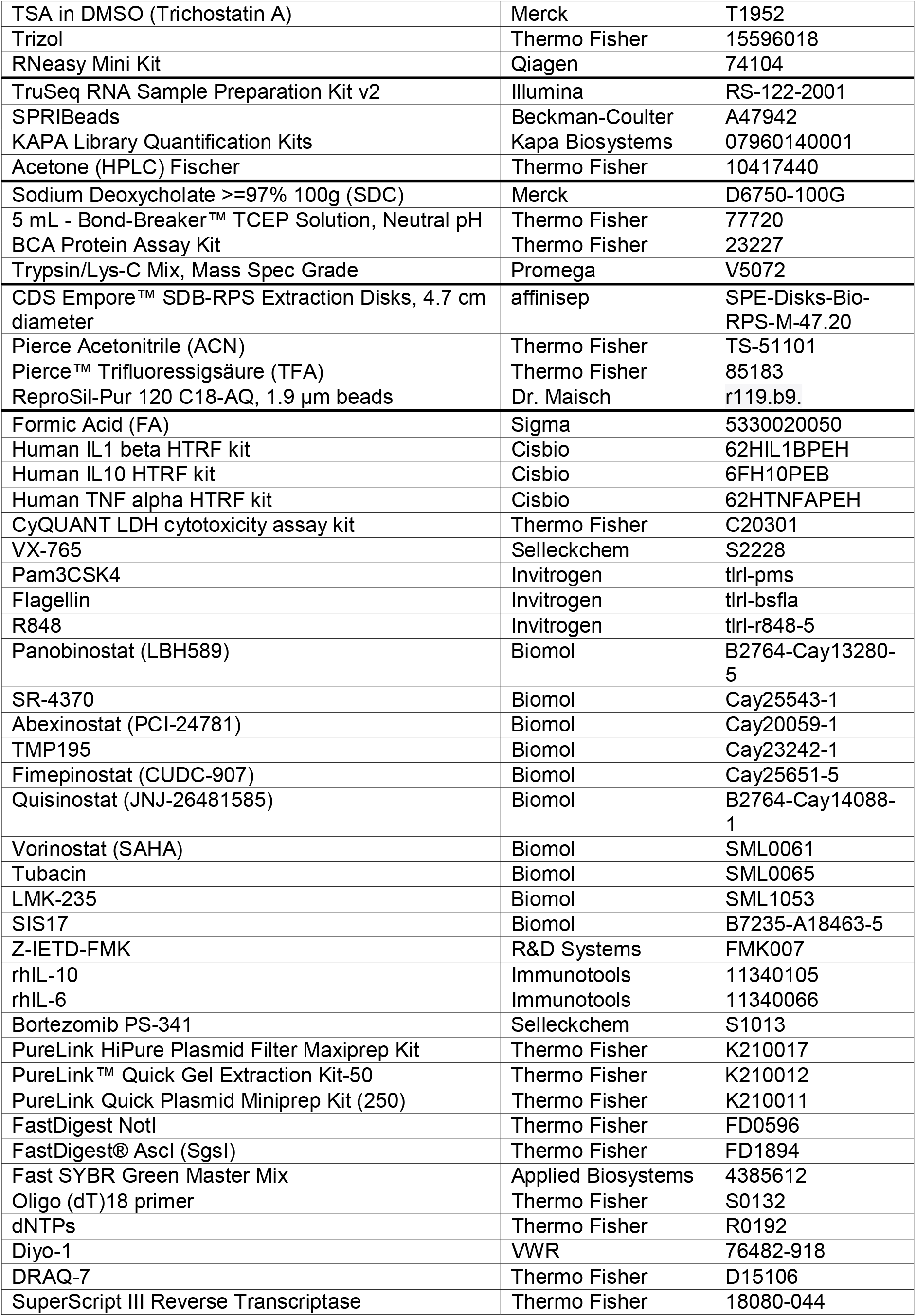
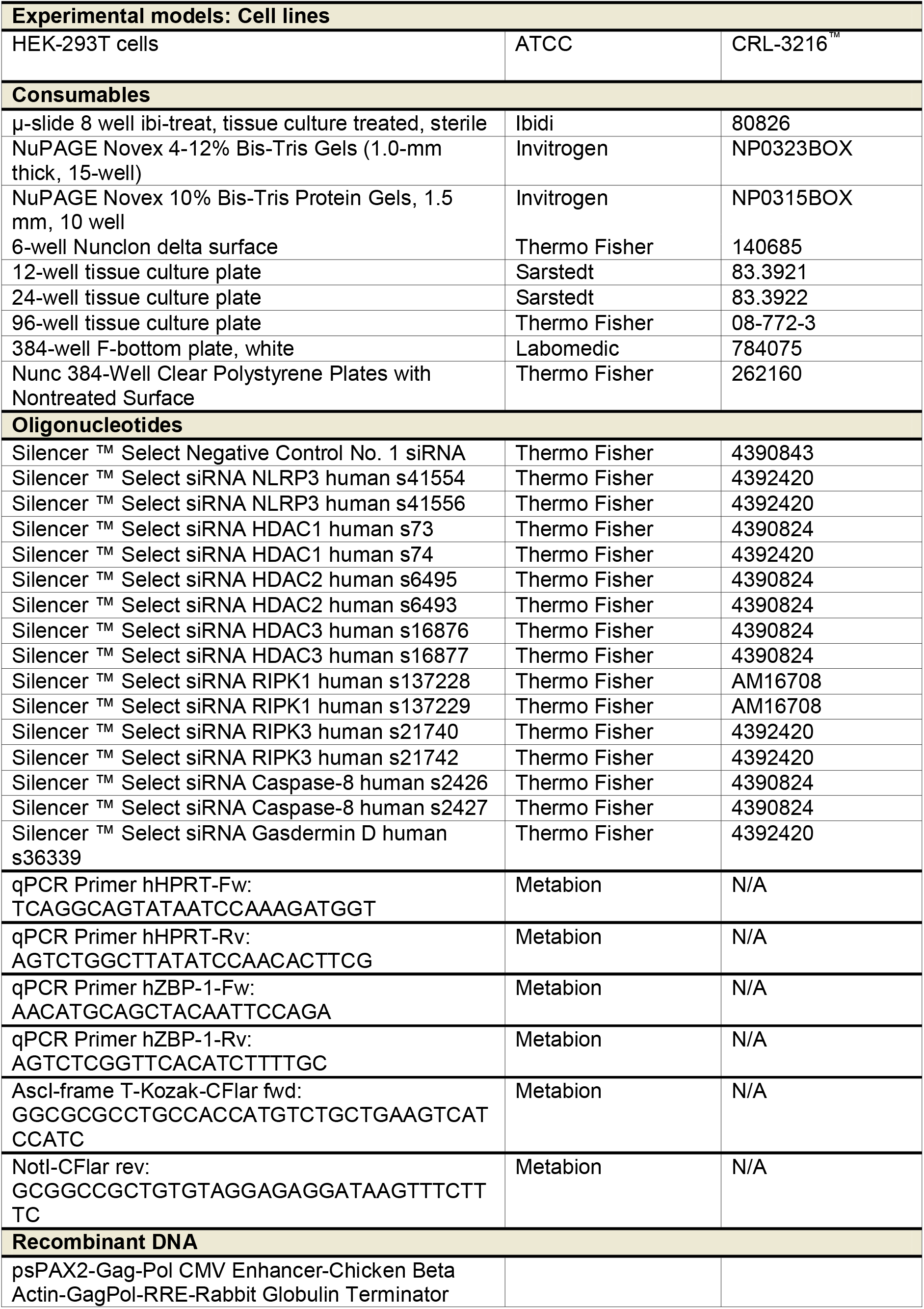
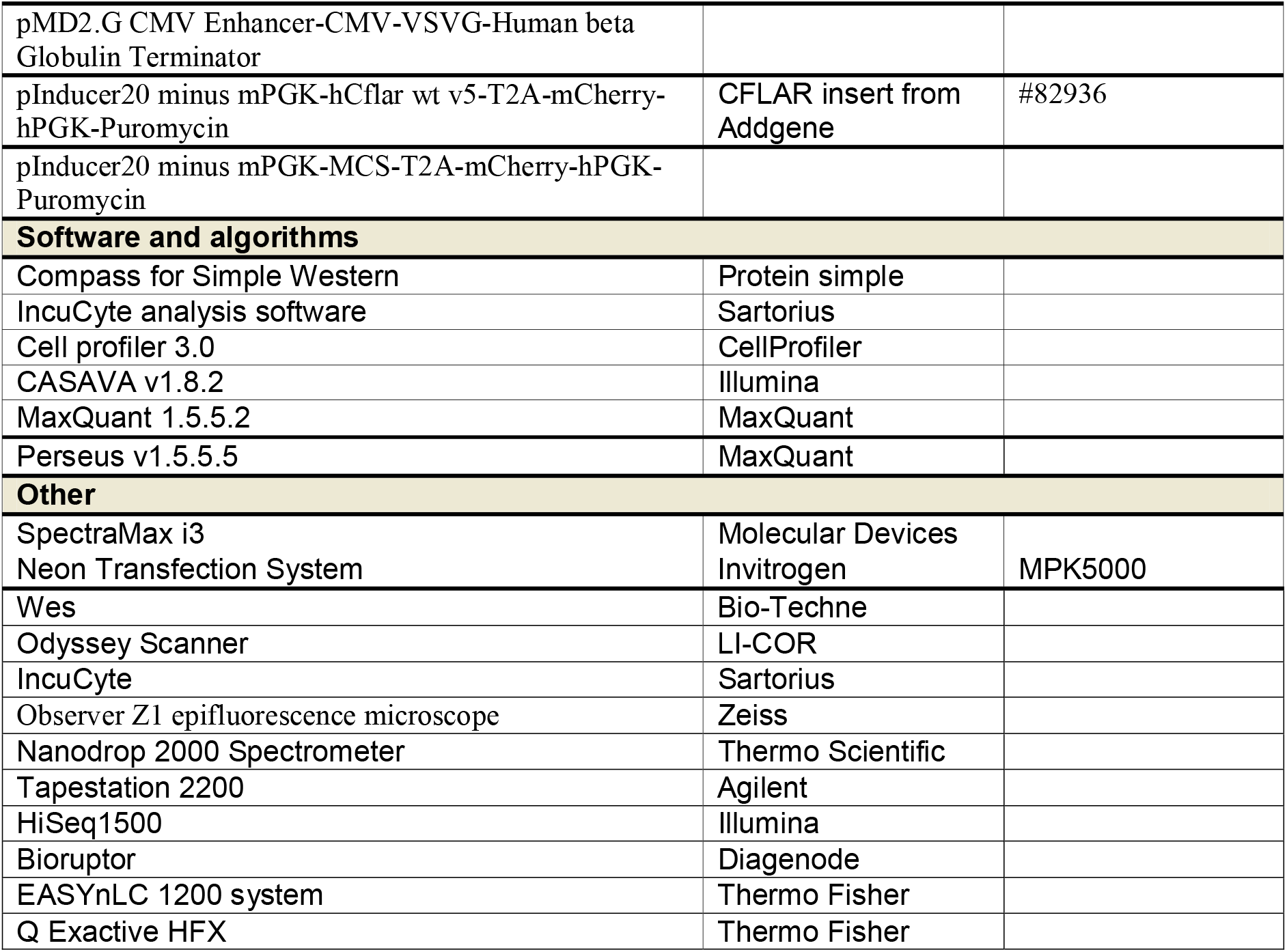

## Data availability

RNA-seq and ChIP-seq data generated for this study have been deposited at the Gene Expression Omnibus (GEO) under accession number GSE248579 (reviewer token: unclakymjrgpxsh)

## Author contributions

E.L, M.M, B.M and W.W conceived the study, W.W, A.D, M.M and E.L designed experiments. W.W, A.D, B.M, M.C, K.P, A.W, N.S, T.W, H.F, A.J, R.S, F.D, S.C and M.M performed experiments. L.L, K.P, C.B, S.S and S.C analysed and provided the visualisation for the RNAseq and ChIPseq data. W.W, A.D, B.M, N.S and M.M performed analysis and visualisation. E.L, M.M, S.C, R.C and F.M supervised the study. M.M wrote the manuscript with input from all authors.

## Acknowledgements

We thank Maximillian Rothe and Romina Kaiser for their technical support and Felix D. Weiss and Daniel Simpson for discussions. pDONR223_CFLAR_WT_V5 was a gift from Jesse Boehm, Matthew Meyerson and David Root (Addgene plasmid #82936; http://n2t.net/addgene:82936; RRID:Addgene_82936). This work was funded in part by the European Union’s Horizon 2020 research and innovation program under grant agreement No. 848146 (To_Aition) (to E.L.), by the Deutsche Forschungsgemeinschaft (DFG, German Research Foundation) under Germany’s Excellence Strategy – EXC2151 – 390873048 (to E.L.), by the DFG SFB1454 - 432325352 (to E.L.), SFB1402 – 414786233 (to E.L.), TRR237 – 369799452 (to E.L.), GRK2168 – 272482170 (to E.L.). S.C. and M.C. were supported by ‘La Caixa Foundation’, the Spanish Ministry of Science and Innovation and NextGeneration-EU (PID2020-117950RA-I00 and RYC2021-033018-I). L.L. was supported by an EU Horizon Europe Marie Skłodowska-Curie grant No.101068212. Furthermore, the studies were supported by the Helmholtz-Gemeinschaft, Zukunftsthema ‘Immunology and Inflammation’ (ZT-0027). RCC was supported by a Biotechnology and Biological Sciences Research Council New Investigator Research Grant (BB/V016741/1).

## Competing interest declaration

E.L. is co-founder of IFM Therapeutics, Odyssey Therapeutics, DiosCure Therapeutics and a Stealth Biotech. R.C.C. is a co-inventor on patent applications for NLRP3 inhibitors, which have been licensed to Inflazome Ltd. and is a consultant for BioAge Labs. The other authors declare no competing interests.

