## Supplementary_figures for "Butyrate and propionate are microbial danger signals that activate the NLRP3-inflammasome in human macrophages in the presence of TLR stimulation"

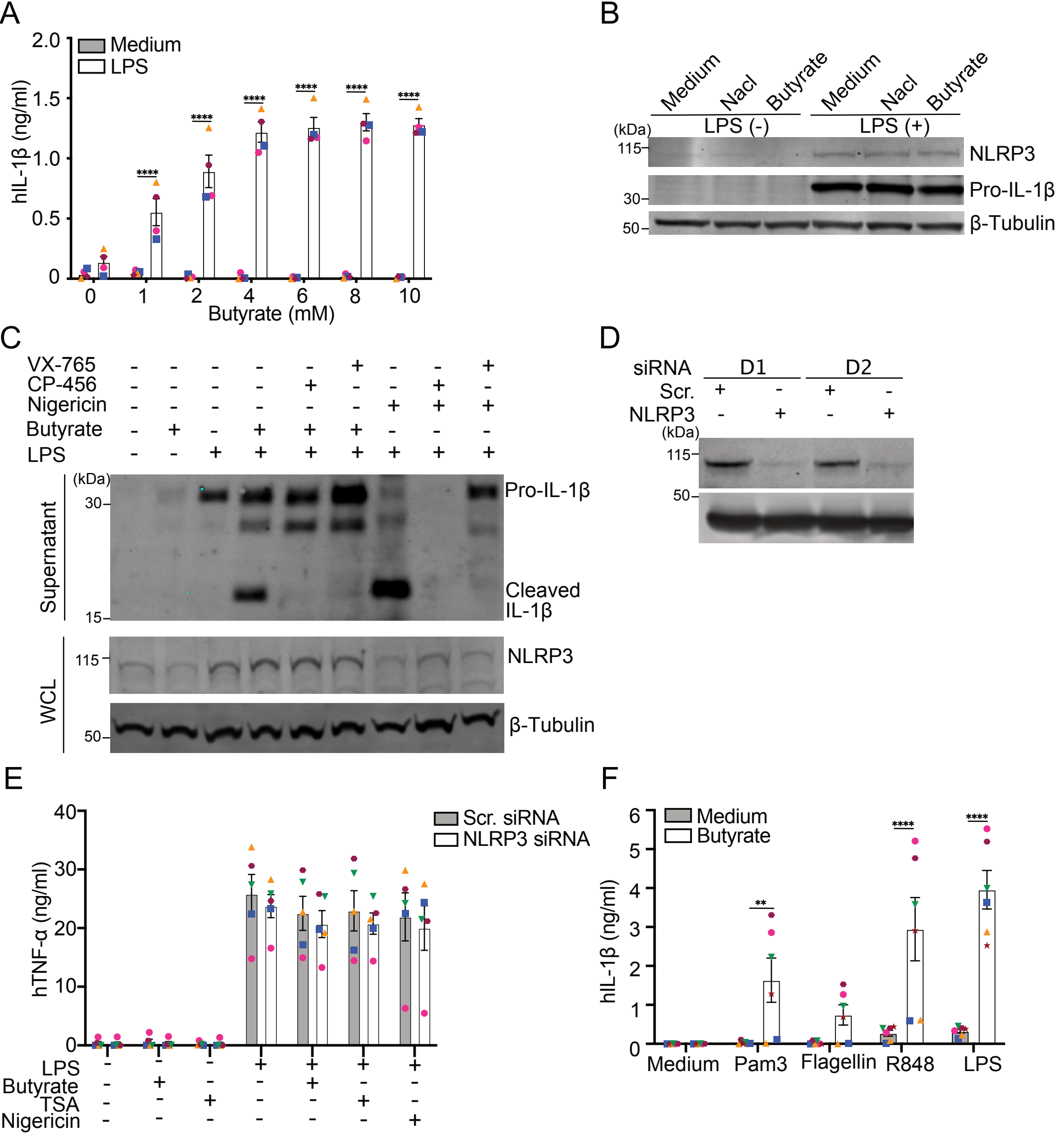
A
Medium
LPS
hIL-1β (ng/ml)
Butyrate (mM)
B
Medium
Nacl
Butyrate
LPS (-)
LPS (+)
(kDa)
NLRP3
Pro-IL-1β
β-Tubulin
C
VX-765
CP-456
Nigericin
Butyrate
LPS
Supernatant
Pro-IL-1β
Cleaved IL-1β
WCL
NLRP3
β-Tubulin
D
siRNA
D1
D2
Scr.
NLRP3
E
Scr. siRNA
NLRP3 siRNA
hTNF-α (ng/ml)
LPS
Butyrate
TSA
Nigericin
F
Medium
Butyrate
hIL-1β (ng/ml)
Medium
Pam3
Flagellin
R848
LPS

Figure S1

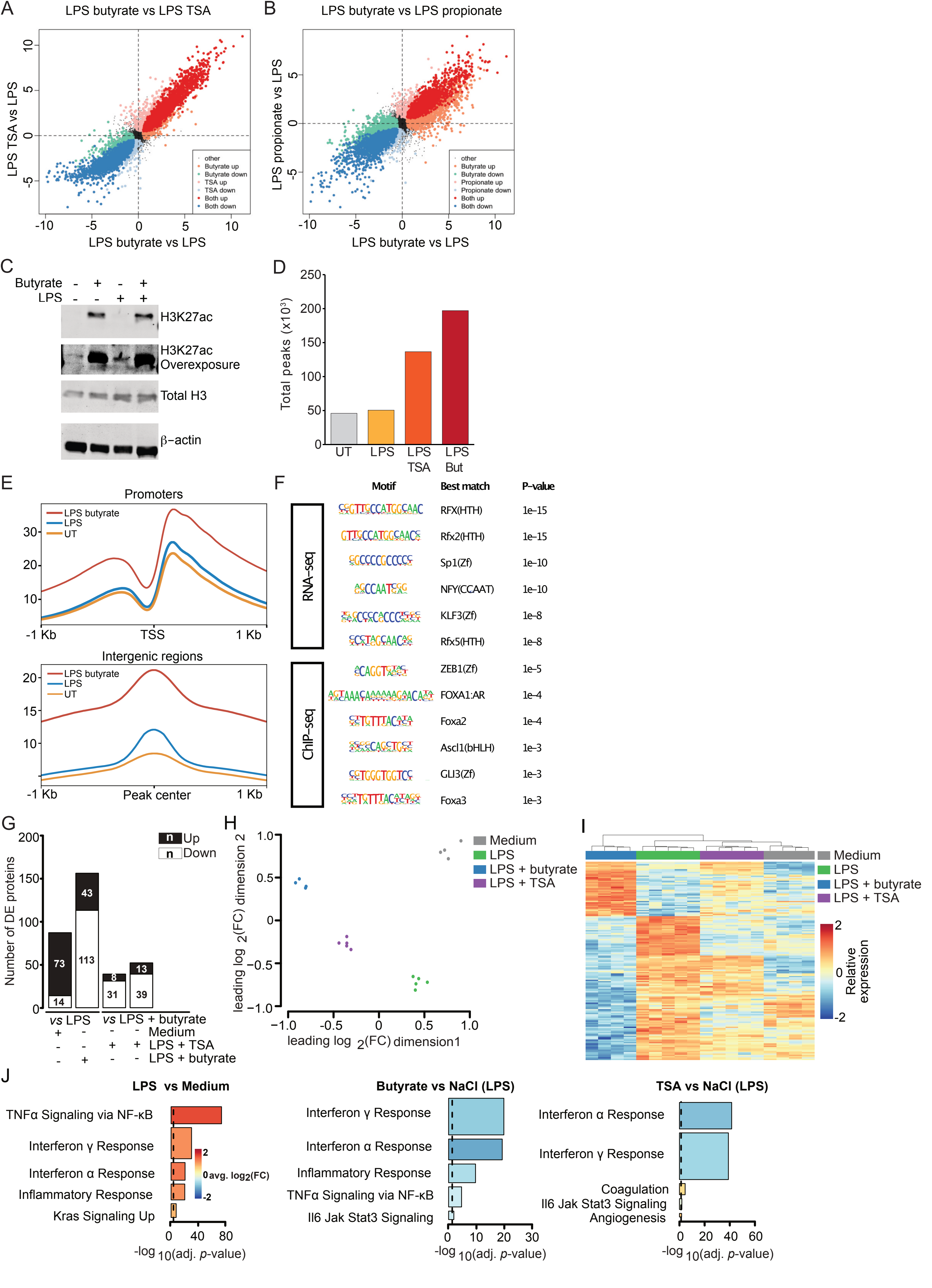
A
LPS butyrate vs LPS TSA
LPS TSA vs LPS
LPS butyrate vs LPS
other
Butyrate up
Butyrate down
TSA up
TSA down
Both up
Both down
B
LPS butyrate vs LPS propionate
LPS propionate vs LPS
LPS butyrate vs LPS
Propionate up
Propionate down
C
Butyrate
LPS
H3K27ac
H3K27ac Overexposure
Total H3
β–actin
D
Total peaks (x10³)
UT
LPS
LPS TSA
LPS But
E
Promoters
LPS butyrate
LPS
UT
-1 Kb
TSS
1 Kb
Intergenic regions
Peak center
F
Motif
Best match
P–value
RNA–seq
ChIP–seq
RFX(HTH) 1e–15
Rfx2(HTH) 1e–15
Sp1(Zf) 1e–10
NFY(CCAAT) 1e–10
KLF3(Zf) 1e–8
Rfx5(HTH) 1e–8
ZEB1(Zf) 1e–5
FOXA1:AR 1e–4
Foxa2 1e–4
Ascl1(bHLH) 1e–3
GLI3(Zf) 1e–3
Foxa3 1e–3
G
Number of DE proteins
Up
Down
vs LPS
vs LPS + butyrate
Medium
LPS + TSA
LPS + butyrate
H
leading log 2(FC) dimension 2
leading log 2(FC) dimension1
Medium
LPS
LPS + butyrate
LPS + TSA
I
Relative expression
J
LPS vs Medium
TNFα Signaling via NF-κB
Interferon γ Response
Interferon α Response
Inflammatory Response
Kras Signaling Up
avg. log2(FC)
-log 10(adj. p-value)
Butyrate vs NaCl (LPS)
Il6 Jak Stat3 Signaling
TSA vs NaCl (LPS)
Coagulation
Angiogenesis

Figure S2

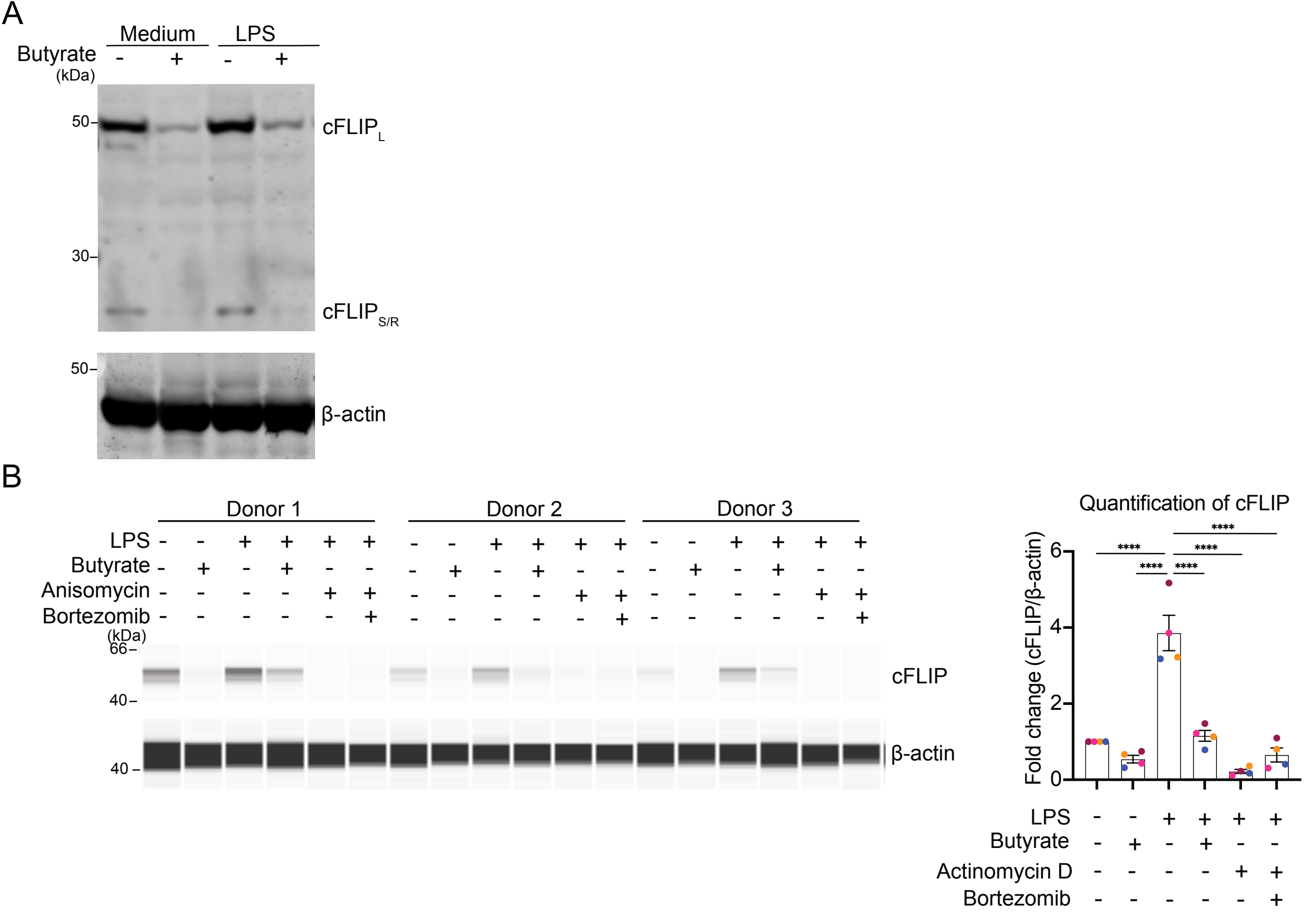
A
Medium
LPS
Butyrate
(kDa)
50
30
cFLIP_L
cFLIP_S/R
β-actin
B
Donor 1
Donor 2
Donor 3
LPS
Butyrate
Anisomycin
Bortezomib
66
40
cFLIP
β-actin
Quantification of cFLIP
Fold change (cFLIP/β-actin)
****
Actinomycin D

Figure S3

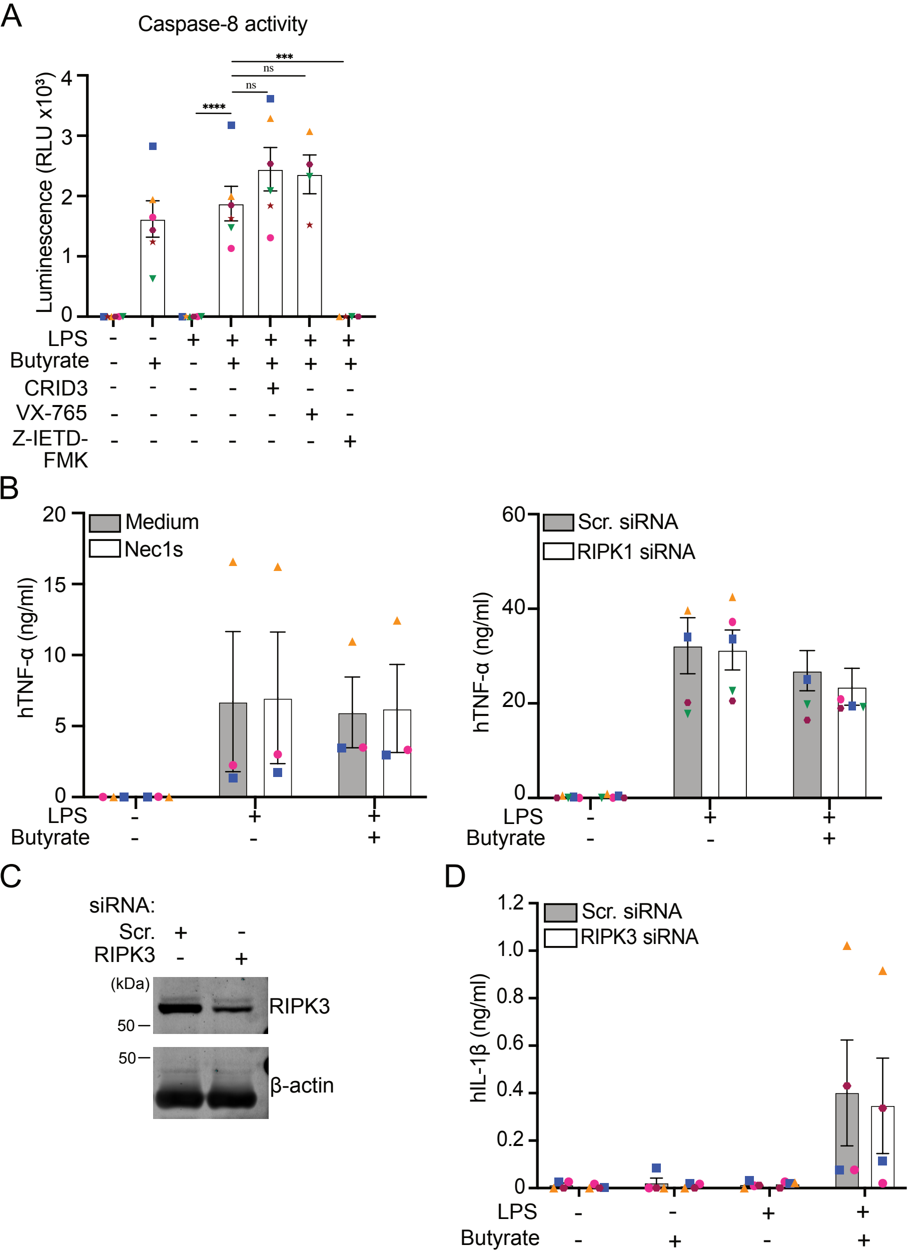
A
Caspase-8 activity
Luminescence (RLU x10³)
****
ns
ns
***
LPS
Butyrate
CRID3
VX-765
Z-IETD-FMK
B
Medium
Nec1s
hTNF-α (ng/ml)
Scr. siRNA
RIPK1 siRNA
hTNF-α (ng/ml)
LPS
Butyrate
C
siRNA:
Scr.
RIPK3
(kDa)
50
RIPK3
50
β-actin
D
Scr. siRNA
RIPK3 siRNA
hIL-1β (ng/ml)
LPS
Butyrate

Figure S4

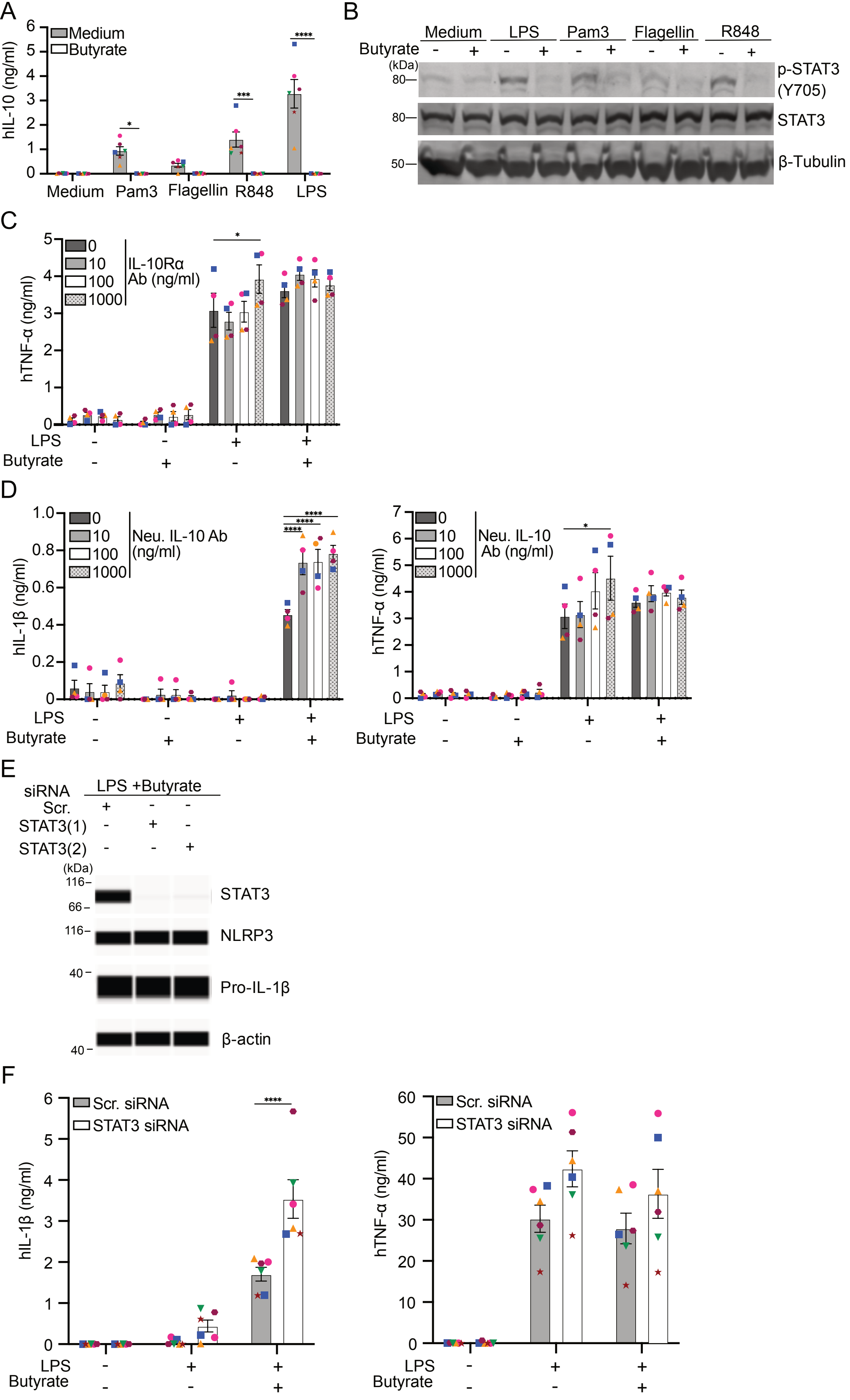
A
Medium
Butyrate
hIL-10 (ng/ml)
Medium
Pam3
Flagellin
R848
LPS
B
Medium
LPS
Pam3
Flagellin
R848
Butyrate
(kDa)
p-STAT3
(Y705)
STAT3
β-Tubulin
C
IL-10Rα
Ab (ng/ml)
hTNF-α (ng/ml)
LPS
Butyrate
D
Neu. IL-10 Ab
(ng/ml)
hIL-1β (ng/ml)
Neu. IL-10
Ab (ng/ml)
hTNF-α (ng/ml)
E
siRNA
LPS +Butyrate
Scr.
STAT3(1)
STAT3(2)
(kDa)
STAT3
NLRP3
Pro-IL-1β
β-actin
F
Scr. siRNA
STAT3 siRNA
hIL-1β (ng/ml)
hTNF-α (ng/ml)
LPS
Butyrate

Figure S5

### Supplementary Figure legends

### Supplementary Figure 1

(A) IL-1β release measured from cell-free supernatants from hMDM treated with various concentrations of butyrate in the absence or presence of LPS (1 ng/ml) for 16 h. Data shown represent 4 independent experiments.

(B) Immunoblot for NLRP3 and pro-IL-1β from hMDM treated with medium, Nacl (10 mM) and butyrate (10 mM) with or without LPS (1 ng/ml) for 16 h. The blots shown are representative of 3 independent experiments. The blots shown are representative of 3 independent experiments.

(C) Immunoblot of cell free supernatants or whole cell lysate (WCL) of hMDM treated with butyrate (10 mM) or Nigericin (10 μM, 1.5 h) with LPS (1 ng/ml) +/- CP-456,773 (2 μM) or VX-765 (40 μM) for 16 h. The blots shown are representative of 2 independent experiments.

(D) Immunoblot of NLRP3 protein from hMDM electroporated with NLRP3 or scrambled siRNA. 2 representative donors shown from 5 independent donors.

(E) TNFα secretion from hMDM electroporated with NLRP3 or scrambled siRNA, then treated with butyrate (10 mM) alone or with LPS (1 ng/ml) for 16 h, TSA (0.5 μM) with LPS (1 ng/ml) for 16 h or nigericin for 1.5 h

(F) IL-1β release from hMDM treated as in Fig. 1F. Data shown represent 4 independent experiments.

Data are represented as mean +/- SEM, unless otherwise indicated, each dot is representative of one donor in all cases, *p<0.05, **p < 0.01, ***p < 0.001, ****p < 0.0001 by Two-way ANOVA with Šídák's multiple comparisons test.

.

### Supplementary Figure 2

FC/FC plot comparing the LPS-Butyrate/LPS condition with either (A) LPS-TSA/LPS or (B) LPS-Propionate/LPS.

(C) Immunoblots of hMDM incubated with LPS and/or butyrate for 6 h then analysed for histone 3 K27 acetylated (H3K27ac), total histone 3 or actin. Representative of 3 independent experiments.

(D) Mean H3K27ac peak calls between replicates for each condition.

(E) Aggregate normalized signal of H3K27ac around TSS and center of intergenic peaks.

(F) Transcription factor motif enrichment analyses for upregulated promoters in Butyrate+LPS vs LPS alone. Motif enrichment was performed at promoters of upregulated genes based on RNA-seq and at the 10% of promoters with the highest H3K27ac increase.

hMDM were treated with either LPS, LPS + butyrate, LPS +TSA or left untreated for 16 hours and the extracted protein was subjected to analysis by mass spectrometry. (G) Barplot depicting the number of differentially expressed (DE) proteins that significantly changed more than 1.2-fold across different comparisons.

(H) Multidimensional scaling (MDS) plot visualising the relationship between the samples, n = 4 or 5.

(I) Heat map depicting relative expression values of proteins that were significantly changed in LPS + butyrate vs LPS comparison, scaled by row.
(J) Gene set enrichment analysis (GSEA) was performed based on the Molecular Signatures Database (MSigDB), using the hallmark gene sets. Plots show the top 5 hallmark gene sets in the following comparisons: LPS vs medium, LPS + butyrate vs LPS and LPS + TSA vs LPS. Bars are coloured by average $\log_2$(FC). Bar width represents the number of genes in the respective gene set. Dashed line indicates adjusted *p*-value threshold.

### Supplementary Figure 3

(A) Immunoblot of cFLIP from hMDM treated with medium, butyrate (10 mM) or LPS (1 ng/ml), alone or in combination, for 16 h. The blots shown are representative of 3 independent experiments.
(B) Immunoblot and quantitation of cFLIP from hMDM were treated with medium, butyrate (10 mM) or LPS (1 ng/ml), alone or in combination, for 6 h pre-incubated with or without Anisomycin (2 µM) or Bortezomib (10 nM) for 0.5 h. Data for 3 donors is presented.

### Supplementary Figure 4

(A) hMDM were treated with medium, butyrate (10 mM) or LPS (1 ng/ml), alone or in combination, for 16 h preincubated with the respective inhibitors 30 min prior to addition of LPS and butyrate (concentrations listed in Figure 3). The activity of the intracellular caspase-8 was assessed using Caspase-Glo® assay kit, luminescence is proportional to caspase-8 activity. Data shown represent 6 independent experiments.
(B) TNFα secretion from hMDM treated as in Fig. 5G and H. Data shown represent 3 (left) or 5 (right) independent experiments.
(C) Immunoblot and (D) IL-1β release from hMDM electroporated with siRNA targeting RIPK3 and incubated with LPS (1 ng/ml) +/- butyrate (10 mM) for 16 h. Data shown represent 4 (D) independent experiments. The blots shown are representative of 3 independent experiments.
Data are represented as mean +/- SEM, unless otherwise indicated, each dot is representative of one donor in all cases, *p<0.05, **p < 0.01, ***p < 0.001, ****p < 0.0001 by either one way ANOVA with Turkey (A) or Two-way ANOVA with Šídák's multiple comparisons test

### Supplementary Figure 5

(A) IL-10 in the cell-free supernatants from hMDM treated with medium or butyrate (10 mM) in the presence of agonists of TLR1/2/6 (Pam3CSK4, 10 ng/ml), TLR4 (LPS, 1 ng/ml), TLR5 (Flagellin, 500 ng/ml), or TLR7/8 (R848, 250 ng/ml) for 16 h. Data shown represent 6 independent experiments.
(B) Immunoblot of phosphorylated STAT3(Y705), total STAT3 and β-Tubulin from hMDM treated with medium, LPS (1 ng/ml), Pam3CSK4 (10 ng/ml), Flagellin (50 ng/ml) or R848 (250 ng/ml) with or without butyrate (10mM) for 16 h. Representative of 3 independent experiments.

(C) TNF-α from cell-free supernatants from hMDM pre-incubated with IL-10Rα blocking antibody for 30 min, then LPS (1 ng/ml) or LPS (1 ng/ml) + butyrate (10 mM) for 16 h. Data shown represent 4 independent donors.
(D) IL-1β and TNF-α measured from cell-free supernatants from hMDM pre-incubated with IL-10 neutralizing antibody for 30 min, then LPS (1 ng/ml) or LPS (1 ng/ml) + butyrate (10 mM) for 16 h. Data shown represent 4 independent donors.
(E) Immunoblots for STAT3, NLRP3, Pro-IL-1β and β-actin from hMDM electroporated with siRNA targeting STAT3 and incubated with LPS (1 ng/ml) +/- butyrate (10 mM) for 16 h 3 days post electroporation. The blots shown are representative of 3 independent experiments.
(F) IL-1β and TNF-α from cell-free supernatants from hMDM electroporated with either scrambled siRNA or STAT3 siRNA then stimulated with medium, LPS (1 ng/ml) or LPS (1 ng/ml) + butyrate (10 mM) for 16 h. Data shown represent 6 independent experiments.

Data are represented as mean +/- SEM, unless otherwise indicated, each dot is representative of one donor in all cases, *p<0.05, **p < 0.01, ***p < 0.001, ****p < 0.0001 by Two-way ANOVA with Šídák's multiple comparisons test.
